# Inferring Bird Migration from Bone Isotopes and Histology: a Fossil-Friendly Methodological Framework

**DOI:** 10.1101/2025.02.08.637244

**Authors:** Anaïs Duhamel, Aurore Canoville, Arnaud Vinçon-Laugier, Julien Joseph, François Fourel, Christophe Lécuyer, Romain Amiot, Antoine Louchart

**Affiliations:** Univ Lyon, Univ Lyon 1, ENSL, CNRS, LGL-TPE, Villeurbanne, France; Museum für Naturkunde, Leibniz-Institut für Evolutions- und Biodiversitätsforschung, 10115 Berlin, Germany; North Carolina Museum of Natural Sciences, 27601 Raleigh NC, USA; Université Claude Bernard Lyon 1, LEHNA UMR 5023, CNRS, ENTPE, F-69622 Villeurbanne, France

**Keywords:** Bone tissues, Bird migration, Fossil birds, Fractionation equation, Oxygen isotopes, Stable isotopes

## Abstract

1. Bird seasonal migration is a remarkable biogeographic phenomenon, yet its deep-time origin(s) and evolutionary history remain poorly understood, with the bird fossil record largely overlooked. This study explores the predictability of bird migratory behaviour from the oxygen isotope composition of their bone apatite phosphate (δ^18^O_p_), a promising approach in this regard because: (i) sedentary and migratory birds tend to occupy distinct climatic niches year-round; (ii) their δ^18^O_p_ values primarily reflect the climate-driven isotopic composition of their drinking water; and (iii) this isotopic signature can persist through fossilisation.
2. Bone tissues were categorised based on their potential to yield spatio-temporally distinct climatic records: *Early Bone Tissues* (EBT), deposited before somatic maturity, and *Late Bone Tissues* (LBT), formed through bone remodelling over the lifespan. The predictability of migratory behaviour was theoretically assessed by modelling tissue-specific δ^18^O_p_ values for thousands of birds across 77 migratory and sedentary species, using tracking and observational data along with a revised phosphate-water fractionation equation. These theoretical results were confronted with data from 11 extant bird species, obtained using a new experimental framework combining histological and isotopic analyses.
3. A significantly positive correlation between δ^18^O_p_ and the proportion of LBT in bone samples was observed both theoretically and experimentally in migratory birds – particularly in long-distance migrants – and is predicted to be virtually absent in sedentary species at temperate latitudes. Migratory birds that died outside their natal sites can also be identified when their δ^18^O_p,EBT_ values fall outside the range observed in local juvenile and sedentary birds.
4. We conclude that bird migratory behaviour can be inferred from the δ^18^O_p_ values of their skeletal remains, provided the individuals migrated across sufficiently contrasting climatic zones. While this approach cannot detect all migratory behaviours – such as short-distance or longitudinal movements – it is unlikely to misclassify sedentary birds as migratory at temperate latitudes. We therefore argue that this approach can be extended to well-preserved fossil bones to infer past bird migratory behaviours, as long as the palaeoclimatic context is carefully considered. More broadly, this study establishes a framework for inferring migratory behaviour in any vertebrate from non-fully remodelled biomineralised remains.

## 1 INTRODUCTION

The evolution of birds has been profoundly shaped by the emergence of one of the most extensive biogeographic phenomena observed among vertebrates: seasonal migrations. Today, billions of birds, representing 20% of all avian species (Somveille, 2016), migrate twice a year between their breeding and wintering grounds, sometimes covering several thousand kilometres. This behaviour is thought to have evolved as an adaptive response to seasonally fluctuating environmental pressures, driven by a complex interplay between abiotic factors (e.g., climate seasonality) and biotic factors (e.g., competition for land and resources, predation and pathogens; Alerstam et al., 2003; Salewski & Bruderer, 2007). However, the deep-time origins and evolutionary history of avian migration – particularly in response to past climate changes (Louchart, 2008) – remain poorly understood. This knowledge would be valuable for deciphering current migratory patterns and anticipating the response of bird species to the ongoing anthropogenic climate and habitat changes.

To date, studies on the origin(s) and evolution(s) of bird migration have largely relied on projecting present-day patterns of biogeography or diversity into the past, through ancestral niche reconstruction (Ponti et al., 2020; Somveille et al., 2020; Dufour et al., 2024) or ancestral state reconstruction approaches (Dufour et al., 2020, 2024). However, this comes with several limitations. First, biotic factors are difficult to assess in deep geological time, and are therefore only partially or not at all considered when inferring past ecological niches. Second, extant life on Earth is a biased sample of all the diversity of life that has existed through geological time (Raup, 1994). Therefore, modelling ancestral states and ancestral ecological niches without integrating fossil evidence significantly reduces inference performance (Finarelli & Flynn, 2006; Meseguer et al., 2015; Lloyd & Slater, 2021). This is particularly true for highly plastic traits such as bird migration, which can switch repeatedly between resident and migratory states across populations and species (Berthold et al., 1992; Helbig, 2003; Bearhop et al., 2005; Zink, 2011; Bensch et al., 2023).

However, whether the avian fossil record can provide insights into past migratory behaviours remains underexplored. Indirect evidence exists at some fossil-rich sites: for certain species, the statistical absence of breeding-related osteological features – such as juvenile specimens (Cheneval, 1989) or medullary bone tissues in females (Matthiesen, 1990) – has been interpreted as evidence of non-breeding presence, suggesting migratory behaviour. Yet such inferences may reflect sampling or preservation biases and are rarely applicable to less rich sites. Alternatively, the biogeographic distribution of fossils has often been misinterpreted as evidence for the spatiotemporal origins of migratory behaviour within lineages, when it only informs the geographic origins of lineages that are migratory today (Bell, 2005). In other words, “fossil birds may indicate where species were present, but not if they migrated”, reinforcing the idea that “migratory behaviour is not recorded in the [avian] fossil record” (Somveille et al., 2020). However, this overlooks the fact that well-preserved fossil material can be vector of palaeoecological information, beyond biogeographic considerations. Notably, chemical elements in hard tissues can preserve their pristine stable isotope compositions during the fossilisation process (Iacumin et al., 1996), and some stable isotopes constitute climatic and/or spatial proxies that appear particularly interesting when it comes to tracking the diverse habitats experienced by migratory animals over their lifetimes.

In terrestrial vertebrates, the stable isotope compositions of soft and hard tissues, such as bones, feathers and teeth, are predominantly influenced by the individual’s intake, including both diet and drinking water. For wild obligate drinkers, the oxygen and hydrogen isotopes in tissues are largely derived from drinking water reservoirs (e.g., rivers, ponds, lakes, snowpacks; Longinelli, 1984). These freshwater reservoirs originate from meteoric waters collected at local or regional scales, whose isotopic compositions correlate positively with air temperature and negatively with precipitation amount, resulting in overall depletion of heavy isotopes with increasing latitude and altitude (Dansgaard, 1964). Relying on stable isotopes that reflect such spatial patterns, protocols have been developed to infer the movements of extant and past vertebrates from the isotopic composition of their biological remains (Hobson & Wassenaar, 2019). These protocols typically rely on tissues that can be easily calibrated over time, such as those with incremental (e.g. teeth, Beard & Johnson, 2000; tusks, Rowe et al., 2024) or seasonal deposition patterns (e.g., feathers, Lott & Smith, 2006; Hobson & Kardynal, 2023). However, these tissues are either absent or poorly preserved in the Cenozoic avian fossil record, which predominantly consists of bone remains. Additionally, most modern birds, with the exception of a few species (e.g., kiwi), reach adult size in less than one year and thus do not exhibit the incremental deposition of bone tissues observed in most other vertebrates (Chinsamy & Elzanowski, 2001).

In many vertebrates, linking stable isotope ratios of bone samples to specific spatio-temporal deposition contexts is complicated by the remodelling of bone tissues throughout life (de Buffrénil & Quilhac, 2021b). In most mammals, bone tissue undergoes complete turnover due to high remodelling rates, and is therefore assumed to reflect a stable isotope value averaged over the individual’s final years of life (e.g., 10 years in humans; Dupras & Schwarcz, 2001; Szostek et al., 2015). As a result, such bones can only inform long-term migratory patterns, and only when compared with tissues that are formed early in life and do not remodel, such as tooth enamel. In birds, however, as in some other vertebrates, some long bones do not remodel as extensively (Starck & Chinsamy, 2002; Currey et al., 2017), allowing certain bone tissues deposited during growth at the breeding site to remain unresorbed over time. Bird bone therefore has the potential to preserve climatic records from distinct spatio-temporal deposition contexts – within tissues that are distinguishable under a light microscope. Properly considered, these temporal records could yield valuable insights into seasonal migration, but this has never been investigated to date.

In this study, we evaluate the predictability of bird migratory behaviour from bone remains using a novel experimental framework that combines stable isotope and histological analyses. Our approach introduces a new classification of bone tissues into *Early Bone Tissues* (EBT), deposited during growth until somatic maturity, and *Late Bone Tissues* (LBT), formed through bone remodelling processes over the lifespan – and thus likely to reflect environmental conditions experienced during migratory stopovers or wintering in migratory species. We hypothesise that differences in the oxygen isotope composition of bone apatite phosphate (δ^18^O_p_) between EBT (δ^18^O_p,EBT_) and LBT (δ^18^O_p,LBT_) can distinguish migratory from sedentary birds, particularly when migratory individuals experienced significant climatic niche shifts between immature and mature life stages.

To test this hypothesis, we developed two complementary predictive approaches. The first, the *Climatic Niche Contrast Test*, classifies as migratory any individual or population showing higher δ^18^O_p_ values in LBT-rich samples relative to EBT-rich samples – a pattern expected in individuals that overwinter in environments warmer than their breeding sites (niche-switching migrants). The second, the *Non-local Origin Test*, classifies as migratory any individual whose EBT isotopic value falls outside the local breeding-season isotopic range, assuming the individual was not born locally and died during a migratory stopover or on the wintering grounds. As a proof of concept, we first assessed the robustness of these approaches using δ^18^O_p_ values simulated across diverse migratory, phenological, physiological and environmental scenarios, and then applied both tests to modern bird specimens with known migratory behaviours and ecological contexts. Finally, we discuss the expected performance of this approach when applied to well-preserved fossil birds, depending on their paleoclimatic and stratigraphic contexts.

## 2 MATERIAL AND METHODS

### 2.1 Experimental framework

Here, we describe a new experimental framework combining δ^18^O_p_ measurements with histological analysis of bone samples, applied as a case study to several migratory and sedentary specimens described in (2.1.1).

#### 2.1.1 Specimens

We selected nine skeletal specimens from five sedentary and four migratory extant bird species, spanning a range of migration routes and timings between Europe and Africa (Fig. S1). Only specimens with a documented deathplace were included, to ensure individuals were not captive during their lifetimes and, in the case of partially migratory species, to confirm they belonged either to strictly migratory or strictly resident populations within the species’ range. All specimens were obtained from the Collections of the Université de Lyon 1 or the Musée des Confluences, with catalog numbers listed in Table S1. Sedentary individuals include *Corvus corone, Corvus frugilegus* (juvenile), *Strix nebulosa* and *Tyto alba*, while migratory individuals include *Ciconia ciconia, Circaetus gallicus, Falco subbuteo, Grus grus* and *Glareola pratincola* (juvenile). As a sodium perborate treatment was historically performed to clean some skeletal remains from the Collections de l’Université de Lyon 1, we tested – and confirmed – that this treatment did not alter the original δ^18^O_p_ values of the bones sampled (protocol and results detailed in Appendix A.1 and Fig. S2).

#### 2.1.2 Sampling protocol

While, ideally, EBT and LBT would be sampled separately, LBT patches are too small and not easily localisable along the bone diaphysis *a priori* without destructive sampling or synchrotron imaging (see 2.1.3, Appendix B and Fig. S3 for detailed histological descriptions of EBT and LBT). We therefore conducted bulk sampling of skeletal elements, assuming each bone sample is a mix of EBT and LBT in proportions *f*_*EBT*_ and *f*_*LBT*_ respectively. The bulk oxygen isotope composition of a sample *(δ*^*18*^*O*_*p, bulk*_) is thus considered a linear combination of its tissue-specific *δ*^*18*^*O*_*p*_ values (*δ*^*18*^*O*_*p,EBT*_ and *δ*^*18*^*O*_*p,LBT*_, equation 1). To better constrain the deposition timing of the sampled tissues, we locally sampled bone diaphyses along cross sections using a Dremel© tool with a diamond-coated circular saw, and paired each bone powder sample from non-juvenile bones with an adjacent thin section to measure the sample-specific *f*_*LBT*_ and *f*_*EBT*_ (2.1.3). All bird species selected (2.1.1) were large enough to yield the minimum 15 mg of bone powder required for the isotopic analysis (2.1.4) within a relatively thin portion of the diaphysis (2-3 mm). The proportion of bone tissue observed on the thin section was thus assumed representative of the associated bone powder sample.

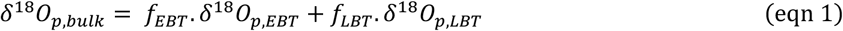

For each individual, we selected a tibiotarsus – known to be relatively rich in remodelled tissues (i.e., LBT) – and an ulna, known to remodel little at the midshaft level (De Margerie et al., 2004; Chinsamy et al., 2014). Ulnae were sampled only once for two main reasons. First, *f*_*LBT*_ was assumed negligible at the midshaft (median 1.1%, Table S1), making the mid-ulnar *δ*^*18*^*O*_*p,bulk*_ value a reliable estimate of the samples’ *δ*^*18*^*O*_*p,EBT*_ value. Second, *δ*^*18*^*O*_*p,EBT*_ was assumed homogeneous within individuals, as it is unaffected by body heterothermy (control developed in Appendix A.2, Fig. S2) and because minor mineralisation timing differences between and within bones of juveniles were considered negligible (Watanabe, 2018; Canoville et al., 2025). Given the limited knowledge about the timing of LBT deposition throughout the year (see 4.1.2), and to maximise the likelihood of sampling LBT patches mineralised outside the breeding season, tibiotarsi were sampled up to eight times along the diaphysis, depending on its length (Fig 1a, Table S1).

**FIGURE 1:**
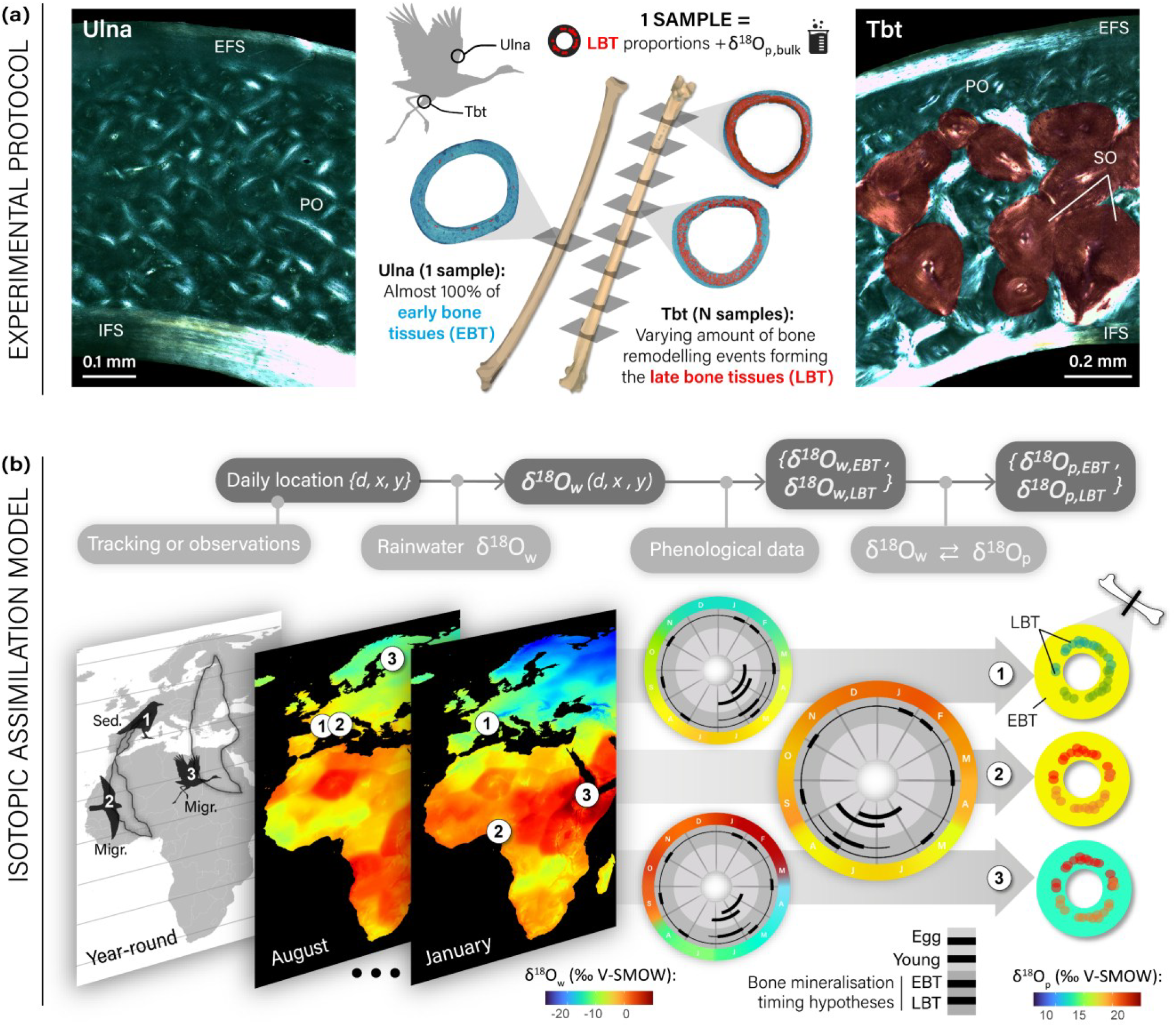
Graphical schematics of the experimental (a) and theoretical (b) protocols deciphering oxygen isotope values in early bone tissue (EBT) and late bone tissues (LBT). (a) Sampling protocol along the ulna and the tibiotarsus (Tbt), and associated thin sections observed under polarized light to quantify EBT (blue) and LBT (red) proportions. Ulna and Tbt thin sections are from *Circaetus gallicus*, and *Ciconia ciconia* respectively. EFS: external fundamental system; IFS: internal fundamental system, PO: primary osteons; SO: secondary osteons. (b) δ^18^O_p_ values modelled in bird bone tissues depending on the hypothetical annual movements of the bird, its experienced δ^18^O_w_ values in precipitation, its species phenological data, and the fractionation equation between δ^18^O_w_ and δ^18^O_p_ in birds. The δ^18^O_w_ values experienced by the birds throughout the year are shown as colour variations around their phenology diagrams. The final outputs are schematic transverse sections, where EBT and LBT patches are coloured according to their corresponding δ^18^O_p_ values.

#### 2.1.3 Thin section histological analysis

To quantify the relative proportions of EBT (*f*_*EBT*_) and LBT (*f*_*LBT*_) within sampled cortical bone, we conducted detailed histological analyses of thin sections under natural and polarized transmitted light using a LEICA DM750P microscope coupled with a LEICA EC3 camera. Following the nomenclature and definitions of de Buffrénil & Quilhac (2021a, 2021b), all patches of secondary osteons were classified as LBT, except for those (rather exceptional) mineralised before somatic maturity due to calcium metabolism imbalances or repair of early microfractures. All remaining bone tissues — including the woven-parallel complex (i.e. woven bone matrix associated with primary osteons) and the internal and external fundamental systems composed of parallel-fibered tissues — were classified as EBT (Fig 1a). These structures were identified based on established diagnostic criteria (Ponton et al., 2007; Chinsamy et al., 2014; Mitchell et al., 2017) detailed and illustrated in Supporting Information (Appendix B; Fig S3). The surface areas occupied by EBT and LBT were measured in pixels using Adobe Photoshop® on composite images of each thin section, reconstructed from a dozen photographs using the “Photomerge” tool. For each thin section, *f*_*LBT*_ was measured directly, and *f*_*EBT*_ was deduced as the remaining proportion of the total cortical surface.

#### 2.1.4 Oxygen isotope analysis of bone apatite phosphate

Bone bioapatite contains two main oxygen-bearing fractions – phosphate (PO_4_^3−^) and carbonate (CO_3_^2−^) – which discriminate oxygen isotopes differently relative to body water (Iacumin et al., 1996). While both reflect water intake (Longinelli, 1984), only the phosphate fraction is currently suitable for modelling (2.2), thanks to the existing fractionation equation between water intake and phosphate apatite in birds (Amiot et al., 2017). No such equation has yet been established for the carbonate fraction. Therefore, we followed a protocol designed to isolate phosphate-bound oxygen from bone samples. Isotopic compositions are reported in the δ notation relative to V-SMOW (Vienna Standard Mean Ocean Water) standard.

The collected 15 to 20 mg samples of bone powders were treated following the same wet chemistry protocol first described by Crowson et al. (1991) and modified by Lécuyer et al. (1993). Five replicates per sample were then prepared by adding 300 to 400 µg of silver phosphate together with 400 µg of pure carbon in silver foil capsules. CO gas formed during the pyrolysis of the capsules performed at 1450 °C using a Elementar vario PYRO cube™ elemental analyser was analysed using an Isoprime™ isotope ratio mass spectrometer. To control both for instrumental drift and for any bias that might occur during the chemistry protocol, δ^18^O_p, bulk_ measurements were calibrated against the phosphorite NBS120c (δ^18^O_p_ = 21.7 ‰ V-SMOW; Lécuyer et al., 1993), the barium sulphate NBS127 (δ^18^O_p_ = 9.3 ‰ V-SMOW; Halas & Szaran, 2001), and an in-house made silver phosphate (Lécuyer et al., 2019) reference materials. The final δ^18^O_p, bulk_ value reported for a sample corresponds to the mean corrected value of the five replicates measured. The mean analytical error across all sample series was estimated at ± 0.22 ‰ V-SMOW.

### 2.2 Theoretical framework

We introduce here a step-by-step theoretical framework (Fig. 1b) to simulate data comparable to those obtained through the experimental approach described above (2.1) – i.e., multiple combinations of bone sample δ^18^O_p_ values (δ^18^O_p,bulk_) and associated histological compositions (*f*_*EBT*_ and *f*_*LBT*_) for each individual – across a broad range of sedentary and migratory bird species. Certain parameters were fixed at the individual level using species- and/or location-specific data available in the literature, while others – poorly documented – were varied systematically as part of a sensitivity analysis (Table 1). Both model implementation and data processing were conducted in R version 4.1.2.

**TABLE 1:** Overview of parameters used to model tissue-specific and bulk δ^18^O_p_ values in bird bones. Parameters with alternative plausible values were explored through sensitivity analyses (see Supplementary Information), with default values used in the main text shown in bold italic. In the “Source” column, “Literature-based” indicates values inferred from empirical data in birds or other vertebrates.

| Output | Parameter | Symbol | Value(s) | Unit | Source |
| --- | --- | --- | --- | --- | --- |
| **Daily $\delta^{18}\text{O}_w$ experienced by the bird** | Calendar day | $d$ | [1,365] | - | - |
| | Daily spatial coordinates (long., lat.) | $x_d, y_d$ | Individual-specific | Decimal ° | Movebank, ebird |
| | Monthly mean $\delta^{18}\text{O}_w$ at position | $\mu_{w,\text{month}}(x_d, y_d)$ | Individual-specific | ‰ V-SMOW | Bowen & Revenaugh, 2003 |
| | Associated interannual SD | $\sigma_{w,\text{month}}(x_d, y_d)$ | | | |
| **$\delta^{18}\text{O}_{w,\text{EBT}}$ (avg. over EBT deposition)** | Egg incubation duration | $D_{\text{inc}}$ | Species-specific (Table S1) | Days | Birds of the World |
| | Laying start and end days | $[d_{\text{ls}}, d_{\text{le}}]$ | Species- and location-specific (Table S1) | - | |
| | Hatching day | $d_{\text{hatch}}$ | $[d_{\text{ls}} + D_{\text{inc}}, d_{\text{le}}]$ | - | - |
| | EBT deposition duration | $D_{\text{growth}}$ | {15, **30**, 45} | Days | Literature-based (2.2.2) |
| | Max. natal dispersal distance | $\omega_{\text{max}}$ | {**0**, 1000} | km | Literature-based (2.2.2) |
| **$\delta^{18}\text{O}_{w,k}$ (avg. over a bone rem. event)** | Secondary osteons deposition duration | $D_R$ | {1, **10**, 30} | Days | Literature-based (2.2.2) |
| | Starting day of the remodelling event | $d_k$ | **[1,365]**, $d_{\text{hatch}} + D_{\text{growth}}$ | - | Literature-based (4.1) |
| **$\delta^{18}\text{O}_p$, tissue ($\delta^{18}\text{O}_p - \delta^{18}\text{O}_w$ regression)** | Regression's slope | $a$ | 0.61 | - | Revised after Amiot et al. (2017) |
| | Regression's intercept | $b$ | 20.97 | ‰ V-SMOW | |
| | Regression's residuals ($\epsilon$) SD | $\sigma_\epsilon$ | {0.50, **1.00**, 1.67} | ‰ V-SMOW | Literature-based (2.2.3) |
| **Bone sample's $\delta^{18}\text{O}_{p,\text{bulk}}$** | Number of bone remodelling events | $R$ | {1, **[1, 5]**} | - | Arbitrary |
| | Proportion of LBT | $f_{\text{LBT}}$ | [0,1] | - | Typical (Table S1) |
| **Sampling scenario** | Number of bone samples | $N_s$ | {5, 7, **9**, 20, 100} | - | - |
| | Number of individuals sampled within the population | $N_i$ | {1, $N_s$} | - | - |

#### 2.2.1 Modelling δ^18^O_w_ values experienced throughout the year

Let *x*_*d*_ and *y*_*d*_ be the longitude and latitude of a bird on calendar day *d* ∈ [1,365], extracted from open-access spatial tracking datasets (2.2.1.1). For each day *d*, the δ^18^O_w_ of local meteoric water was modelled using global monthly maps of mean δ^18^O_w_ values *(µ*_*w,month*_*(x*_*d*_, *y*_*d*_)) and associated interannual standard deviations (*σ*_*w,month*_*(x*_*d*_, *y*_*d*_)) (2.2.1.2). Importantly, *σ*_*w,month*_ describes variability across years, not day-to-day fluctuations. Consequently, a single residual ε_*w,month*_ was drawn per location per month from a normal distribution (equations 2, 3). Daily temperature and precipitation values were simulated similarly.

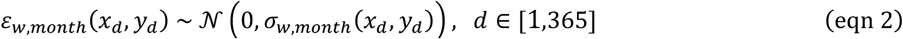

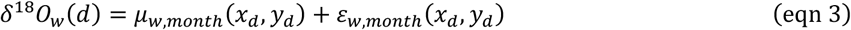

##### 2.2.1.1 Year-round location data

To account for as much variability in extant bird migratory patterns as possible – i.e. intra- or inter-specific variability in migration timing or routes – we downloaded from the Movebank database (https://www.movebank.org/, accessed in February 2024) the tracking data of 184 individuals from 33 migratory species (Fig. S5, see Table S1 and Data Source for associated references). These raw tracking data were subsampled to retain only one spatial coordinate per day over a year. Missing daily locations, often due to interrupted tracking during long stopovers, were filled by the spatial coordinates of the latest available location. Non-overland locations – for which there is no correspondence on land climatic and isotopic maps (2.2.1.2) – were replaced by the spatial coordinates of the latest overland location, assuming that the δ^18^O_w_ values recorded in bone tissues during these marine flyovers are those derived from the latest land water supply.

To account for as much variability in extant sedentary birds habitats and phenologies as possible, we drew several living locations for 44 sedentary bird species across Europe, western Asia and Africa from the revised 2023 eBird Basic Dataset (http://www.ebird.org, accessed in February 2024; Sullivan et al., 2009). To improve spatial coverage across each species’ range, the probability to draw an observation was weighted by the relative frequency of observations per country (Fig. S4). Each draw from a species-specific database was assumed to be the year-round living location for an individual of that species.

##### 2.2.1.2 Climatic and isotopic data maps

The raster grids of global monthly δ^18^O values of precipitation (δ^18^O_w_) and associated standard deviations were extracted from the Waterisotope portal (http://waterisotopes.org, Bowen & Revenaugh, 2003). The raster files of global monthly air temperatures, precipitation amount and associated standard deviations averaged from 1970 to 2000 were extracted from the WorldClim 2.1 database (Fick & Hijmans, 2017). To avoid potential missing values for coastal spatial coordinates (e.g., wetlands), the land-water boundaries of all these raster grids were slightly dilated by averaging the nearest land pixels values using the *focal* function of the *raster* package (Hijmans et al., 2015).

#### 2.2.2 Modelling δ^18^O_w_ values experienced over EBT and LBT deposition

##### 2.2.2.1 EBT deposition

EBT deposition was assumed to occur linearly from hatching (day *d*_*H*_) to somatic maturity, over a duration *D*_*EBT*_. Accordingly, δ^18^O_w_ experienced during EBT deposition was modelled as the average of local daily values over this period:

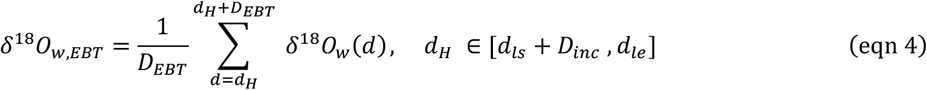

The pre-hatching bone fraction of EBT was neglected for three reasons: (i) chicks have very low body mass at hatching compared to adults; (ii) pre-hatching tissues of long bones are largely resorbed during post-hatching growth (de Buffrénil, 2021); and (iii) maternal-to-embryo water transfer may have introduced additional δ^18^O_w_ discrimination.

Somatic maturity was assumed to coincide with the attainment of adult body size, which aligns with fledging in most species (Ricklefs, 1968, 1979; Carrier & Auriemma, 1992). Based on observed fledging timing across birds species of comparable body mass (Carrier & Auriemma, 1992; Cooney et al., 2020), *D*_*EBT*_ was set to values varying within {15,30,45} days. For each individual, plausible hatching dates (*d*_*H*_) were estimated by shifting species- and location-specific laying periods (from *d*_*ls*_ to *d*_*le*_) by the incubation duration of the species (*D*_*inc*_), using data from *Birds of the World* (https://birdsoftheworld.org/bow/home, Billerman et al., 2022; Table S1).

In migratory species, EBT deposition was restricted to the period spent in the breeding area. Breeding locations were identified by segmenting year-round movements into steps marked by > ± 2° shifts in latitude or longitude. The breeding step was defined as the longest step occurring within [*d*_*ls*_, *d*_*le*_ + *D*_*EBT*_] period.

Sedentary birds often establish their territory several kilometres away from their natal site (i.e., natal dispersal), most commonly within 20 km (Paradis et al., 1998; Claramunt, 2021; Fandos et al., 2023), with rare occurrences reaching several hundred kilometres (e.g., up to ~650 km in *Strix nebulosa*, Solheim & Stefansson, 2016). Accordingly, we drew natal sites such that their distances from the adult territory approximately follows a uniform distribution between 0 and 1000 km (*ω*_*max*_). Equations 2 and 3 were then applied to these natal sites instead of the positions showed in Fig. S4. For migratory species, natal dispersal was considered negligible relative to seasonal movement.

##### 2.2.2.2 LBT deposition

LBT result from bone remodelling events cumulated over the lifetime, each forming patches of secondary osteons after resorption of underlying tissues (de Buffrénil et al., 2021). The time needed for secondary osteons deposition (*D*_*R*_) is poorly documented in birds, but is known to vary with body size and metabolic rate (de Buffrénil & Quilhac, 2021b). In cortical bone, deposition can last a single day in rats and up to 60-120 days in humans (Vignery & Baron, 1980). Here, *D*_*R*_ was therefore varied across simulations using values of 1, 10, or 30 days. Similarly, little is known about the deposition timing of LBT over the bird’s lifespan (4.1.2). Bone remodelling events were thus either modelled as: (i) randomly distributed across the year, with starting dates drawn uniformly from [1,365], or (ii) occurring exclusively post-breeding on the breeding ground, with a minimum starting date of *d*_*H*_ + *D*_*EBT*_. This latter scenario represents a biologically plausible “worst-case” scenario (discussed in 4.1.2) in which LBT would not be deposited either on the wintering grounds or at migratory stopovers. δ^18^O_w_ experienced during the deposition of a single bone remodelling event *k*, with deposition starting on a date *d*_*k*_, was modelled as:

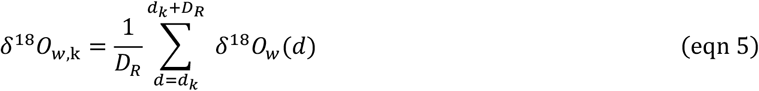

#### 2.2.3 Modelling tissue-specific δ^18^O_p_ values

The rainwater oxygen undergoes several isotopic fractionation steps before being incorporated into bone tissues (Luz et al., 1984). As a result, δ^18^O_w_ and δ^18^O_p_ tend to be linearly correlated. Such a fractionation equation was introduced for birds by Amiot et al. (2017), based on broiler chickens. However, it relied on an Ordinary Least Squares (OLS) regression that minimised residuals only for the dependent variable δ^18^O_w_, which is not suitable for estimating δ^18^O_p_ from δ^18^O_w_. We therefore re-fitted the original dataset from Amiot et al. (2017) using Reduced Major Axis (RMA) regression, which treats both variables symmetrically and is therefore appropriate for predicting either δ^18^O_p_ from δ^18^O_w_ *or vice versa*. This was performed using *lmodel2* R package (Legendre, 2018). Furthermore, Amiot et al. (2017)’s δ^18^O_w_ estimates were based on local annual precipitation averages, whereas the broiler chickens sampled were juveniles and mostly hatched during the season of lowest local δ^18^O_w_ values. To addressed this potential bias, we recalculated δ^18^O_w_ estimates as the 30-day post-hatching average of local δ^18^O_w_ values, and re-fit the RMA regression. We further calibrated this equation using our dataset of juvenile and sedentary birds using the 30-day post-hatching average of local δ^18^O_w_ (Appendix C; Fig. S6). Thus, δ^18^O_w,EBT_ and δ^18^O_w,k_ values were converted into tissue-specific δ^18^O_p_ values using the following oxygen isotope fractionation equation between bone phosphate and rain water (equation 6).

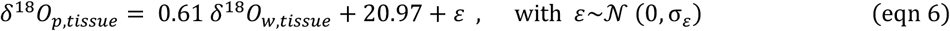

A normally distributed residual term (ε) was added to each δ^18^O_p, tissue_ estimate, reflecting hidden factors (e.g., water source heterogeneity, metabolic rate or physiology; 4.1.4) assumed to remain stable over the short timescales of EBT deposition and bone remodelling events. Because equation 6 was derived from a taxonomically broad dataset including both wild and domesticated birds, its residual standard deviation (σ_ε_= 1.62 ‰) likely overestimates the variability expected within a single individual. To better reflect realistic biological variation, we also performed simulations using σ_ε_values of 1‰ – a more typical within-species estimate based on studies in terrestrial vertebrates (e.g., North American deer; Cormie et al., 1994) – and 0.5‰, representing an optimistic estimate for intra-individual variation.

#### 2.2.4 Modelling a bone samples’ δ^18^O_p,bulk_ values

Since LBT represents cumulative bone remodelling throughout an individual’s life (de Buffrénil et al., 2021), the δ^18^O_p,LBT_ value of a bone sample was modelled as the average of *R* distinct bone remodelling events (equation 7). *R* was either fixed at 1 or allowed to increase proportionally with *f*_*LBT*_, up to a maximum of five events.

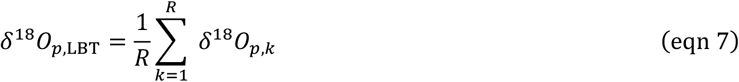

As outlined in Section 2.1.2, bone samples may contain both LBT and EBT fractions, in proportions *f*_*LBT*_ and *f*_*EBT*_ *= 1 - f*_*LBT*_, respectively. These proportions vary depending on the sampled element and species-specific life history traits (discussed in 4.1.2). To reflect this variability, *f*_*LBT*_ was randomly drawn from a uniform distribution over [0,1]. The resulting bulk δ^18^O_p_ value of a bone sample was then calculated as:

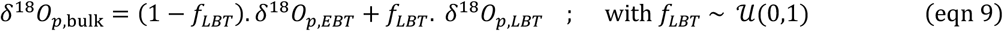

### 2.3 Statistical analyses

Here we introduce two complementary statistical frameworks designed to detect migratory behaviour from histologically informed δ^18^O_p_ data.

#### 2.3.1 Testing for climatic niche contrasts

The *Climatic Niche Contrast Test* was developed to detect migratory behaviour by comparing δ^18^O_p_ values between EBT and LBT. The underlying hypothesis is that migratory birds – especially those wintering in warmer climates – should exhibit higher δ^18^O_p_ values in LBT than in EBT, whereas sedentary species inhabiting regions with cooler year-round average temperatures relative to the growth season may show the opposite pattern.

In theory, δ^18^O_p,LBT_ could be isolated from a sample’s bulk measurement (δ^18^O_p,bulk_) using equation 1 and an estimated δ^18^O_p,EBT_ value from another sample with negligible *f*_*LBT*_ (assuming δ^18^O_p,EBT_ is homogenous within a single individual; 2.1.2). However, this approach introduces substantial uncertainty– especially when *f*_*LBT*_ is low – because of error propagation (Table S1). Alternatively, we inferred migratory behaviour from the correlation between δ^18^O_p, bulk_ and *f*_*LBT*_. As this relationship is not necessarily linear, we used Spearman’s rank correlation coefficient (*ρ*) to quantify its direction and strength.

The correlation was assessed across *N*_*S*_ samples (i.e., pairs of δ^18^O_p, bulk_ and *f*_*LBT*_ values), obtained either from a single individual or from multiple individuals within a population (i.e., conspecifics from the same locality; in fossil assemblages, multiple individuals may need to be sampled to achieve adequate sample size, 4.2.2). Two independent classification rules were applied in separate tests: individuals or populations showing a significantly positive correlation (ρ>0 and p-value<0.05) were classified as migratory (“*ρ>0\**” decision rule), while those with a significantly negative correlation (*ρ*>0 and p-value<0.05) were classified as sedentary (“*ρ<0\**” decision rules).

Tests performances were evaluated theoretically in terms of sensitivity (true positive rate) and specificity (true negative rate) across 100 replicates, each consisting of *N*_*S*_ randomly sampled δ^18^O_p, bulk_ - *f*_*LBT*_ pairs (2.2). Each sampling scenario – either within individuals (*N*_*I*_ *= 1* individual sampled) or within populations (*N*_*I*_ *= N*_*S*_ individuals sampled) – was tested across a range of sample sizes (*N*_*S*_ = 5, 7, 9, 20 or 100) and model parameters (Table 1). Individuals within a population were modelled with similar annual movements (except natal dispersal in sedentary species) but potentially differing hatching timings.

#### 2.3.2. Testing for non-local origin

The *Non-local Origin Test* was developed to identify any individual whose EBT isotopic values fall outside the local isotopic range, under the assumption that some migratory individual may have died during a migratory stopover or on their wintering grounds. A species was classified as migratory if any EBT-rich sample had a δ^18^O_p_ value falling outside the 95% confidence interval of the theoretical or empirical δ^18^O_p,EBT_ distribution for local birds. Empirical distributions were derived from ulnar mid-shaft samples and tibiotarsus samples with over 80% EBT content. Theoretical distributions were based on modelled δ^18^O_p,EBT_ values for local sedentary birds observed in the wild (2.2).

## 3 RESULTS

### 3.1 Contrasting δ^18^O_p, EBT_ and δ^18^O_p, LBT_ values

#### 3.1.1 Theoretical tissue-specific climatic pattern

To evaluate whether migratory birds can be distinguished from sedentary birds based on their EBT and LBT δ^18^O_p_ values, it is first essential to assess how environmental δ^18^O in precipitation (δ^18^O_w_) varies across life cycles and regions, and how reliably these variations are reflected in bone tissues δ^18^O_p_ values.

##### 3.1.1.1 δ^18^O_w_ variations over life cycles

In both temperate sedentary (25°–67° latitude) and migratory birds, chick growth is highly seasonal, predominantly occurring between April and August (Fig. 2a). In temperate latitudes, the growth season coincides with the onset of warmer temperatures and an increase in δ^18^O_w_ of local precipitation (Fig. 2a). Consequently, most temperate sedentary individuals (91.8%) experience lower average δ^18^O_w_ values year-round than during their growth period (Fig. 3a1-a2). The few individuals showing the opposite trend in average (8.2%) generally appear as population-level outliers, as this occurs in only 2.6% of populations (Fig. 3a1-a2). In migratory birds that travel short latitudinal distances (0°–15°) or migrating longitudinally (e.g., Richard’s Pipit, *Anthus richardi*, Fig. S4), non-breeding climates tend to resemble those of their breeding zones. Their life-cycle δ^18^O_w_ values therefore show similar or less pronounced seasonal contrasts than those of temperate sedentary birds (Figs. 2b, 3a1-a2, 4b). By contrast, long-distance migrants covering broader latitudinal distances (15°–102° latitude) experience warmer – and occasionally drier – conditions during non-breeding periods (Fig. 2b). For species such as the Common Ringed Plover (*Charadrius hiaticula*) or the Caspian Tern (*Hydroprogne caspia*), temperature differences between breeding and non-breeding periods can reach up to ~35°C. This results in higher δ^18^O_w_ during non-breeding season (Figs. 2b, 3a1-a2, 4b). A similar trend was observed in some sedentary individuals and populations from tropical regions (~25° to −35° latitude; Fig. 3-4a), where hatching is less seasonal and likely occurs during the late rainy season, when δ^18^O_w_ is lower (Fig. 2a).

**FIGURE 2:**
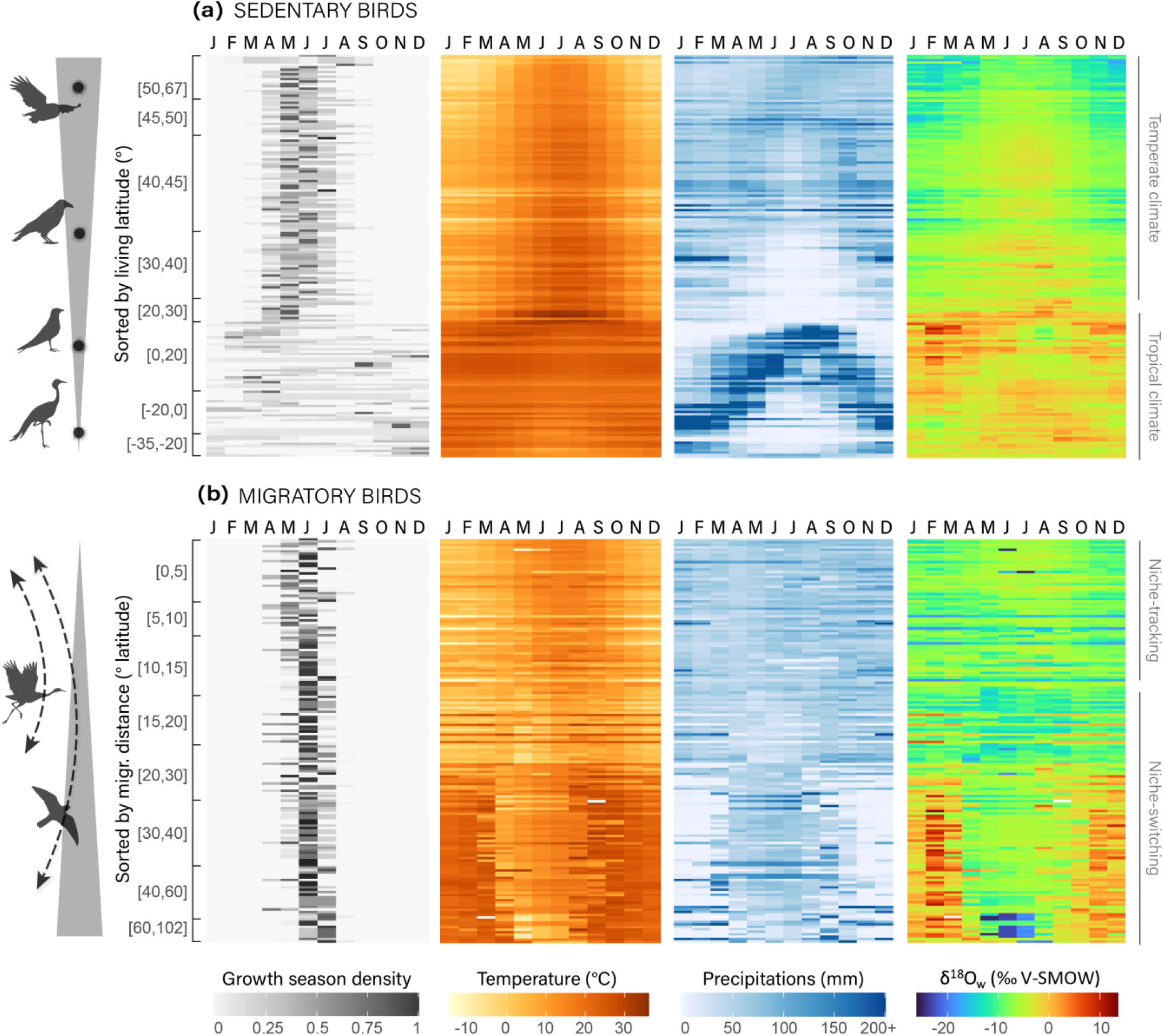
Mean monthly climatic variables (temperatures, precipitation amount and precipitation δ^18^O_w_) theoretically experienced by sedentary or migratory populations throughout a year (from January to December), based on their year-round spatial coordinates (Fig. S4-S5), compared to the local growth season density of their species. Each row represents a population (i.e., individuals of the same species sharing similar year-round coordinates), with sedentary populations sorted by living latitude (a), and migratory populations sorted by migratory distance (in degrees latitude) (b).

**FIGURE 3:**
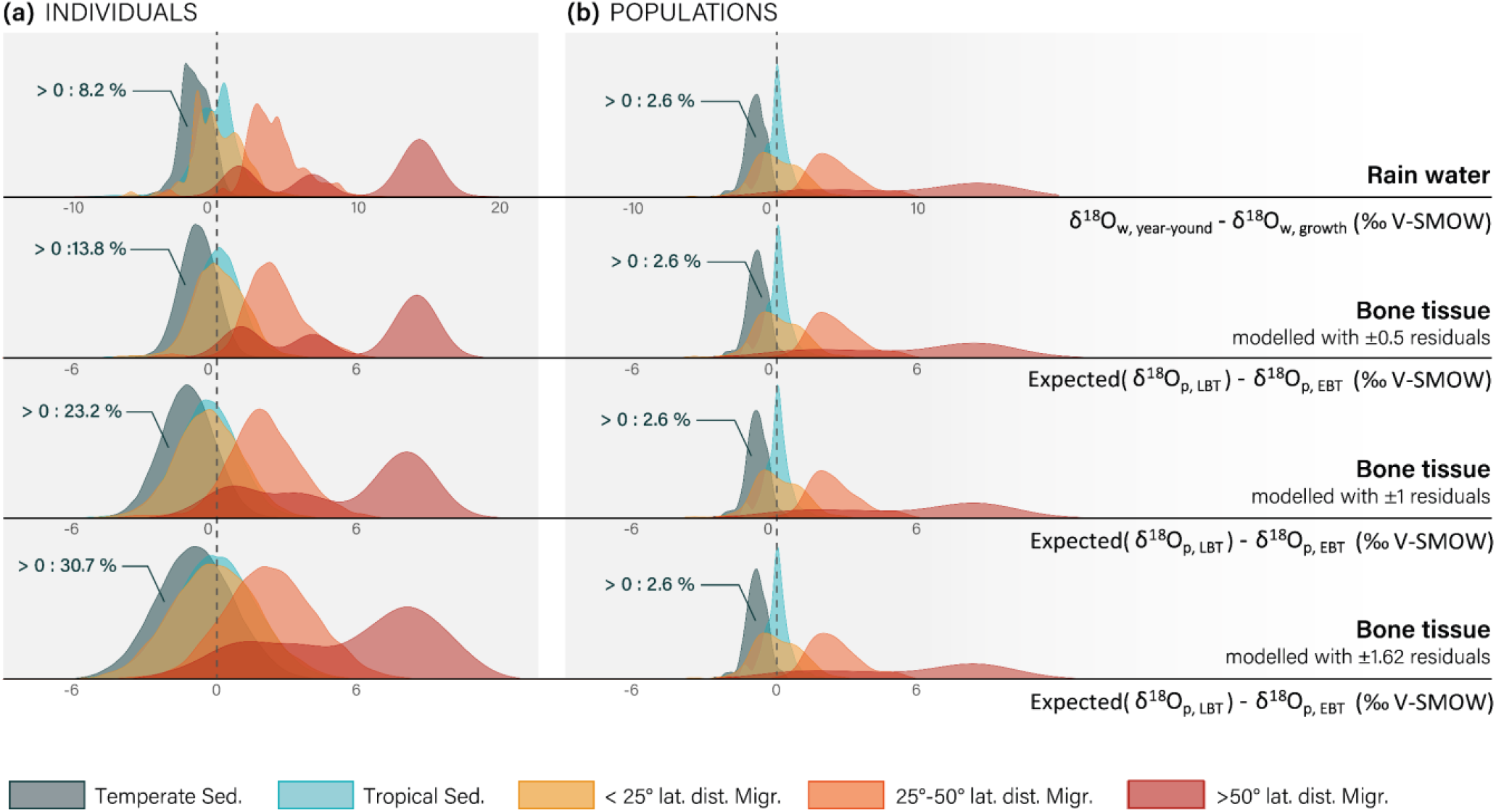
Theoretical contrasts between δ^18^O_w_ values of year-round and growth-period precipitation, and the resulting δ^18^O_p_ contrasts expected in bone tissues deposited during these periods (LBT and EBT, respectively), depending on the type of migratory behaviour. These contrasts are shown for (a) individuals and (b) populations (i.e., averaged over individuals of the same species sharing similar year-round locations but with different hatching timings). δ^18^O_p_ values were estimated using equation 6, with normally distributed error equal to either ± 0.5 ‰, 1 ‰ or 1.62 ‰, and other model parameters set to their default values (see Table 1). The expected value of δ^18^O_p, LBT_ represents the long-term isotopic composition integrated over potential remodelling events.

**FIGURE 4:**
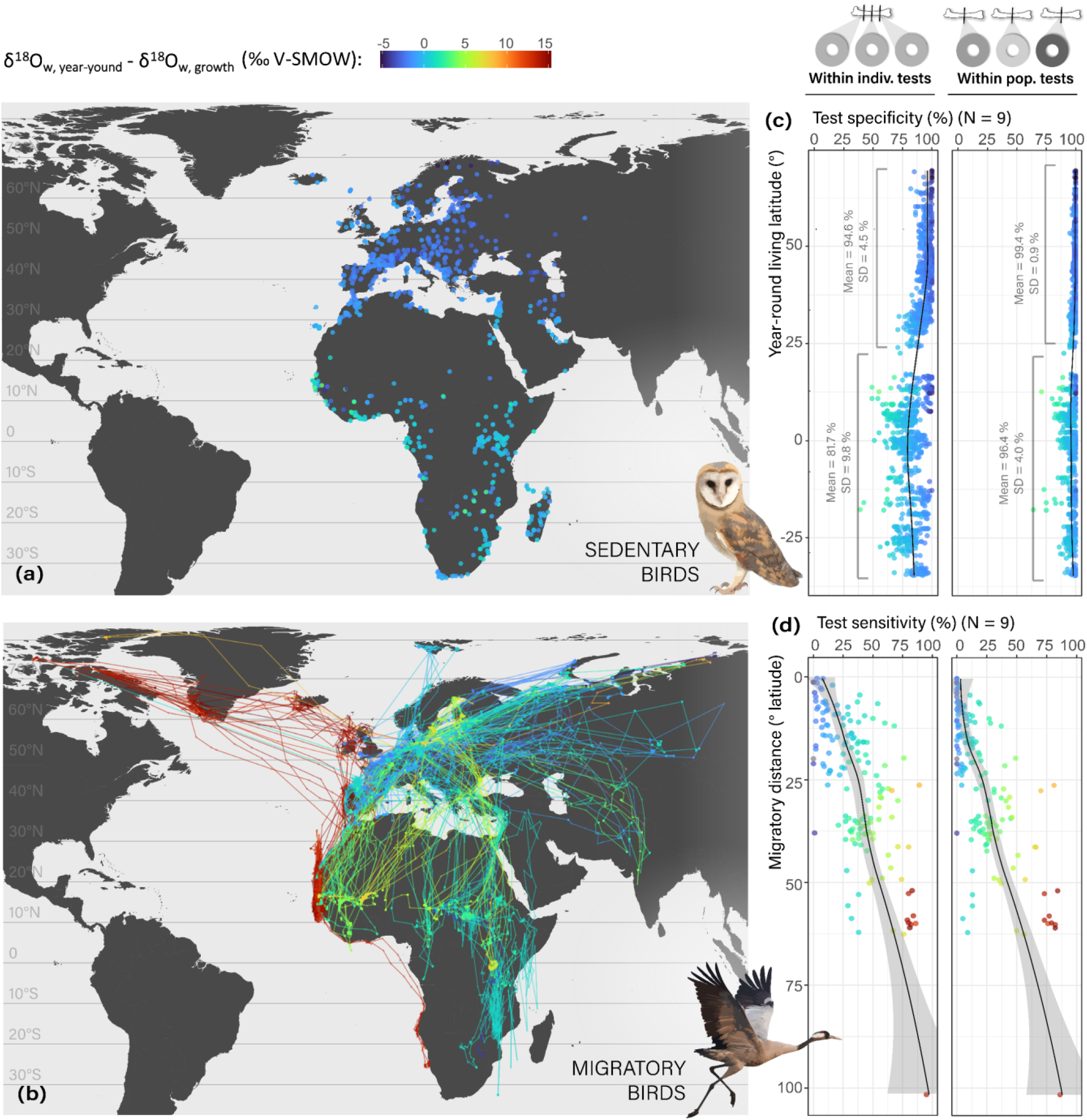
Year-round locations of sedentary birds (a) and tracking coordinates of migratory birds (b) coloured according to the population’s mean difference between δ^18^O values of year-round and growth-period rainwater. Theoretical specificity (c) and sensitivity (d) of the “Climatic Niche Contrast Test”, using the “*ρ*>0 *” decision rule: i.e., individuals or populations are classified as migratory (“positive” outcome) when they exhibit significantly positive correlations between *f*_*LBT*_ and δ^18^O_p, bulk_ (Spearman’s *ρ*>0 and p-value<0.05). This decision rule was applied to samples of size *N*=9, drawn either from repeated measurements within a single individual (fixed δ^18^O_p, EBT_ values; left column plots) or from individuals within the same population (varying δ^18^O_p, EBT_ values; right column plots). All other model parameters were fixed at their default values (see Table 1). Sensitivity (proportion of true positives) and specificity (1 – proportion of false positives) were computed from 100 *N*-sized samples, for each individual or population. Specificity was plotted for sedentary individuals or populations against their living latitude, while sensitivity was plotted for migratory individuals or populations against their migratory distance. Observations are coloured following the same colour gradient as (a) and (b).

##### 3.1.1.2 δ^18^O_p_ contrasts between bone tissues

We next examined tissue-specific δ^18^O_p_ values, first modelled using default parameters (Table 1 or Fig. 5), which notably assume random deposition of LBT throughout the year and no natal dispersal. Reflecting the life-cycle variations in δ^18^O_w_ values described above (3.1.1.1), we note, on average (Fig. 3): (i) lower or weakly contrasted δ^18^O_p_ values in LBT than in EBT for both temperate sedentary birds and longitudinal or short-distance migrants (< 25° latitudinal distance); (ii) reduced contrasts in sedentary birds from tropical climates; and (iii) higher δ^18^O_p_ values in LBT than in EBT for long-distance migrants, with contrast magnitude increasing with migratory distance.

**FIGURE 5:**
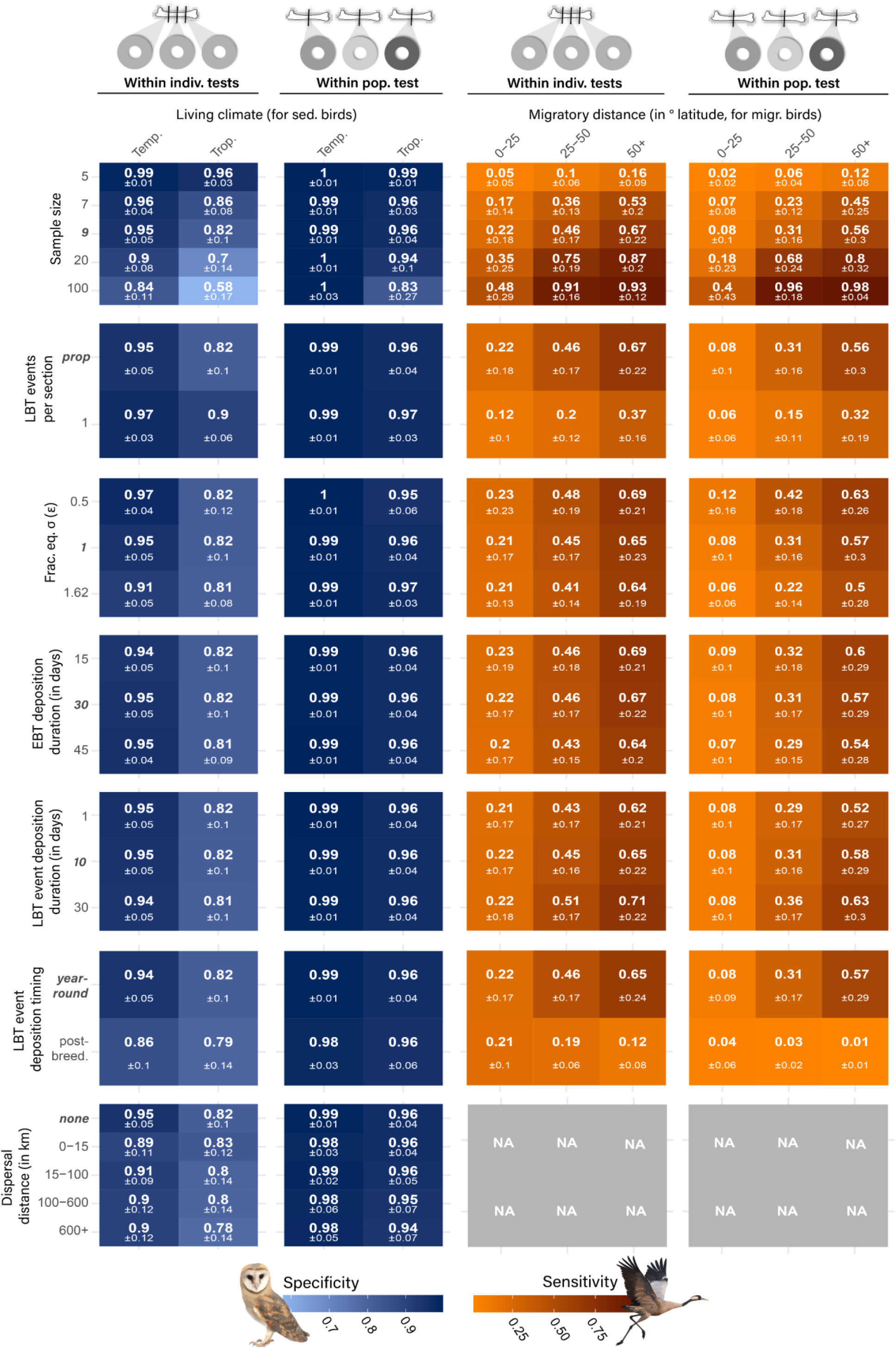
Sensitivity analysis of the “Climatic Niche Contrast Test”, using the “*ρ*>0 *” decision rule, conducted by varying model parameters one at a time while all others were held fixed at their default values (indicated in *italic bold*). The decision rule was applied to samples drawn either from repeated measurements within a single individual (fixed δ^18^O_p, EBT_ values) or from individuals within the same population (varying δ^18^O_p, EBT_ values). Sensitivity (proportion of true positives) and specificity (1 – proportion of false positives) were computed from 100 *N*-sized samples for each individual or population. Specificity is plotted as the mean ± SD for sedentary individuals or populations, grouped by living climate (temperate vs. tropical), while sensitivity is plotted as the mean ± SD for migratory individuals or populations, grouped by migratory distance.

Because of residuals added to the linear relationship between δ^18^O_w_ and δ^18^O_p_ (representing random noise or other unmeasured variability, εin equation 6), positive differences between δ^18^O_p, LBT_ and δ^18^O_p, EBT_ values appeared in 13.8% of temperate sedentary individuals with σ_ε_= ± 0.5‰, increasing to 23.2% with σ_ε_= ± 1‰ and up to 30.7% with σ_ε_= ± 1.62‰ (Fig. 3a). However, these positive contrasts remained negligible at the population level, occurring in fewer than 2.6% of populations regardless of residual magnitude (Fig. 3b).

The contrasts described above persisted – with slightly greater variance – when sedentary birds were modelled as dispersing in any direction up to 1000 km from their natal site (Fig. S8 & Fig. S9). However, δ^18^O_p, LBT_ - δ^18^O_p, EBT_ contrasts vanished for both migratory and sedentary birds when LBT deposition was restricted to the breeding ground immediately post-breeding (Fig. S8).

#### 3.1.2 Predictability of migratory behaviour in case of bulk sampling

Since we do not have direct access to tissue-specific δ^18^O_p_ values in practice, we instead measure bulk δ^18^O_p_ values from bone samples containing varying proportions of LBT and EBT (2.1.2, 2.3.1). Below, we test whether the correlation between the δ^18^O_p, bulk_ and *f*_*LBT*_, calculated from a set of samples taken either from a single individual or from multiple individuals within the same population, enables reliable classification of individuals or populations as migratory or sedentary.

##### 3.1.2.1 In function of sampling strategy

We first vary sampling strategy (individual vs. population; see 2.2.4) and sample size (*N*_*S*_) with all non-sampling-related model parameters set to their default values (see Table 1 or Fig. 5).

Regardless of the sampling strategy and sample size, significantly positive correlations (i.e., *ρ*>0 and p-value<0.05) between *f*_*LBT*_ and δ^18^O_p, bulk_ were observed in both migratory and tropical sedentary species (Figs. S10). In temperate sedentary species, such correlations were found in up to 16.4% of individuals (increasing with *N*_*S*_) but became negligible at the population level as *N*_*S*_ increased (e.g., less than 0.6% of N_S_-sized samples when *N*≥9; Fig. S10-S11).

Classifying individuals as migratory based on significantly positive correlations between *f*_*LBT*_ and δ^18^O_p, bulk_ (“*ρ*>0*” decision rule) therefore resulted in a notable false positive rate among tropical sedentary species, with a specificity of 81.7%, when *N*_*S*_=9 (Figs. 4c-S11). In temperate sedentary individuals, the test yielded high average specificity for small sample sizes (94.6% when *N*_*S*_=9), but specificity declined with larger sample sizes, dropping to 83.6% when *N*_*S*_=100. However, applying the same decision rule at the population level substantially reduced misclassification of temperate sedentary populations as migratory, achieving over 99% specificity across all tested sample sizes, with only a slight reduction of sensitivity. For migratory individuals and populations with positive contrasts between LBT and EBT – i.e., mostly long-distance migrants – sensitivity increased with *N*_*S*_, reaching ~100% for most species when *N*_*S*_*≥*20 (Fig. S11).

Both sedentary birds from temperate climates and longitudinal or short-distance migrants tend to exhibit significantly negative correlations (i.e., *ρ*>0 and p-value<0.05) between *f*_*LBT*_ and δ^18^O_p, bulk_ (Figs. 4a-S10). Consequently, regardless of sample size and sampling strategy, using the “ρ<0 *” decision rule to identify birds as sedentary resulted in low specificity, with over 50% of samplings leading to misclassification of short-distance migratory birds as sedentary (Fig. S12b).

##### 3.1.2.1 Sensitivity analysis

To assess the robustness of these results under varying model assumptions, we conducted a sensitivity analysis of the *Climatic Niche Contrast Test*, using the “ρ>0 *” decision rule. Model parameters were varied one at a time, while all others were held fixed at their default values (Table. 1, Fig. 5).

First, for both sampling scenarios (individual- or population-level), assuming that only a single remodelling event is captured per thin section (R = 1) led to a noticeable drop in sensitivity. This is expected, as the likelihood of capturing LBT deposited during the wintering period decreases under this assumption. Conversely, variation in the durations of EBT and LBT deposition (*D*_*EBT*_ and *D*_*R*_, respectively) had no impact on test specificity or sensitivity in either scenario.

As expected, when LBT deposition is restricted to the breeding ground immediately post-chick growth, test sensitivity dramatically dropped due to highly reduced contrasts between δ^18^O_p, LBT_ and δ^18^O_p, EBT_ values. While this represents a worst-case scenario (discussed in 4.1.3), population-level tests still retained very high specificity.

In both individual- and population-level tests, increasing the stochasticity of δ^18^O_p_ estimates – i.e., the strength of random noise or other unmeasured variability in the linear relationship between δ^18^O_w_ and δ^18^O_p_ (σ_ε_in equation 6) – significantly reduced sensitivity. However, even under the most pessimistic scenario (σ_ε_= 1.62 ‰), the test remained moderately sensitive for birds migrating over more than 50° of latitude (~64% of individuals and ~50% of populations detected as migratory) and highly specific at the population level, with fewer than 1% of temperate sedentary birds misclassified as migratory.

Finally, greater natal dispersal in sedentary birds slightly reduced specificity at the individual level. This effect, however, was mitigated at the population level, where isotopic signals from random dispersal directions tend to average out (Fig. S9a).

#### 3.1.3 Application case

As a consistency check of the model and an illustration of the statistical approach described above, we measured δ^18^O_p_ values and associated *f*_*LBT*_ values (except for juveniles) in three adult sedentary individuals, two juveniles, and four migratory individuals (2.1.1). Using juveniles and sedentary individuals as controls, whose lifelong location are well constrained, we found that δ^18^O_p_ values measured in both LBT and EBT generally matched theoretical expectations (Fig. S7). Among migratory birds, only the Common Crane (*Grus grus*) exhibited a significant positive correlation between δ^18^O_p, bulk_ and *f*_*LBT*_ (ρ = 0.73, p-value = 3.11.10^−2^; Table 2). For sedentary species, only the Great Gray Owl (*Strix nebulosa*) showed a significant negative correlation (ρ = −0.93, p-value = 2.94.10^−4^). Importantly, no sedentary individuals were misclassified as migratory using the “ρ>0 *” decision rule, and no migratory individuals were misclassified as sedentary using the “ρ<0 *” decision rule.

**TABLE 2:** For each adult bird sampled, description of the correlation between the percentage of LBT (100.*f*_*LBT*_) and the δ^18^O_p,bulk_ of the bird’s samples. The Spearman’s ρ and the p-value of this relationship are to be considered together with the SD of both variables and with the sample size (N). Statistically significant Spearman’s rank correlations at the 95% confidence level are indicated by a “**”. For each species, the minimum and maximum migratory distances between breeding and wintering latitudes were estimated from eBird data maps (Fig. S1; Sullivan et al., 2009).

| Specimen’s species | Migr. dist. (° lat.) | Sample size ($N_s$) | $\sigma(\delta^{18}\text{O}_{\text{p, bulk}})$ (‰ V-SMOW) | $\sigma(100.f_{\text{LBT}})$ (%) | Min ($100.f_{\text{LBT}}$) (%) | Max ($100.f_{\text{LBT}}$) (%) | Spearman’s $\rho$ | p-value |
| --- | --- | --- | --- | --- | --- | --- | --- | --- |
| *Tyto alba* | 0 | 7 | 0.1680 | 17.6336 | 0.0000 | 52.0389 | 0.2523 | $5.8524 \cdot 10^{-1}$ |
| *Corvus corone* | 0 | 7 | 0.4432 | 10.2933 | 3.4415 | 33.6281 | 0.3928 | $3.9563 \cdot 10^{-1}$ |
| *Strix nebulosa* | 0 | 9 | 0.7528 | 21.4539 | 0.0000 | 60.3346 | -0.9289 | **$2.9452 \cdot 10^{-4}$**<br>** |
| *Ciconia ciconia* | [15-92] | 9 | 0.4297 | 21.0098 | 0.0000 | 64.7845 | -0.5000 | $1.7766 \cdot 10^{-1}$ |
| *Falco subbuteo* | [20-100] | 6 | 0.4800 | 11.4274 | 0.2715 | 25.7973 | -0.7143 | $1.3611 \cdot 10^{-1}$ |
| *Circaetus gallicus* | [15-62] | 9 | 0.6260 | 16.5966 | 0.0000 | 41.2122 | 0.3529 | $3.1510 \cdot 10^{-1}$ |
| *Grus grus* | [15-55] | 9 | 0.5888 | 21.8825 | 1.4817 | 65.5689 | 0.7333 | **$3.1123 \cdot 10^{-2}$**<br>** |

### 3.2 Locally contrasting δ^18^O_p,EBT_ values

Finally, we applied the *Non-Local Origin Test* to assess whether migratory birds that did not hatch on-site but originated from climatically distinct regions could be distinguished from local birds. The test was performed using both empirical and theoretical δ^18^O_p,EBT_ data for two localities: Southern France and Ivory Coast. Theoretical δ^18^O_p,EBT_ distributions showed lower variance under the temperate climate of Southern France than under the tropical climate of Ivory Coast, and the empirical distributions showed less variance than their theoretical counterparts (Fig 6). *G. grus* exhibited a δ^18^O_p,EBT_ value (13.6‰ V-SMOW) outside the 95% confidence intervals of both control distributions. This value closely matched theoretical expectations for northern breeding areas, such as Scandinavia (Fig. 6a-c), consistent with the species’ documented breeding range (Fig. S1). In contrast, the Short-toed Snake Eagle (*Circaetus gallicus*) and the Eurasian Hobby (*Falco subbuteo*) exhibited δ^18^O_p,EBT_ values within the local distributions, suggesting they were either born at their place of death (consistent with their breeding range) or climatically similar regions. Lastly, while *C. ciconia* cannot have been born in Ivory Coast in the wild given its species distribution (Fig. S1), it could not be classified as non-local with 95% confidence, as its δ^18^O_p,EBT_ value fell within the theoretical distribution for its death site (Fig. 6b). This can be explained by the breeding season δ^18^O_w_ values across the documented breeding range of *C. ciconia* (e.g., southern France, Fig. 6c, Fig. S1), which largely resemble those observed in the Ivory Coast.

**FIGURE 6:**
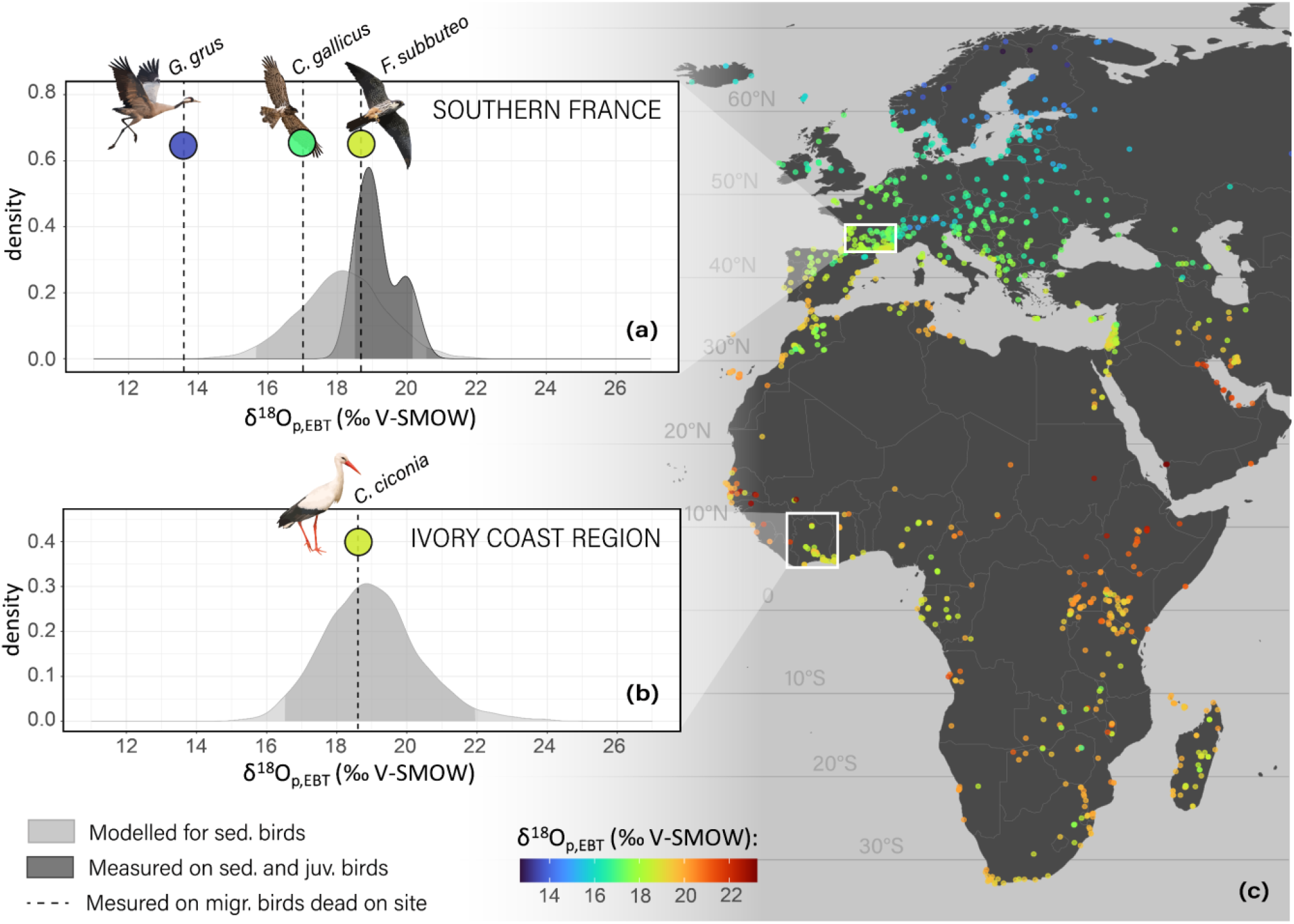
Among migratory birds whose remains were found in either Southern France (a) or in Ivory Coast (b), identification of individuals with δ^18^O_p,EBT_ values falling outside the 95% confidence interval of the theoretical and/or empirical δ^18^O_p,EBT_ distributions of local birds (sedentary and/or juveniles, see 2.3.2). The 95% confidence interval has a darker shade within each distribution. The dots in (c) illustrate the locations of each modelled population, coloured following its mean δ^18^O_p,EBT_ value.

## 4 DISCUSSION

To our knowledge, this study is the first to investigate the predictability of bird migratory behaviour using stable isotope ratios in bone tissues, signatures that can persist through fossilisation. To this end, we developed a protocol that combines δ^18^O_p_ measurements with histological analyses distinguishing bone tissues deposited before somatic maturity from those deposited throughout life through bone remodelling (early bone tissue, EBT, vs. late bone tissue, LBT). As a proof-of-concept, we demonstrated through a comprehensive simulation framework that migratory behaviour can be confidently inferred from such data when birds experienced sufficient seasonal climatic contrasts, and we further illustrated the experimental feasibility of this approach. Below, we discuss the assumptions underlying this method and its applicability to deep-time material.

### 4.1 Inferring migratory behaviour from bone isotopes: scope and best practises

#### 4.1.1 Two complementary tests for detecting migratory patterns

Climatic signatures, both geographic and seasonal, are intrinsically embedded in rainwater δ^18^O_w_ values (Dansgaard, 1964). We demonstrated that two complementary tests allow the prediction of specific migratory behaviours based on the δ^18^O_w_ values of water metabolised by birds during their EBT and LBT deposition periods (i.e., growth and lifetime, respectively), even when accounting for monthly and interannual variations in rainwater δ^18^O_w_ values.

##### (i) Climatic Niche Contrast Test

By contrasting the mean δ^18^O_p_ of bone tissues deposited over the lifetime (LBT) with those deposited during growth (EBT), we emphasised that niche-switching migratory birds, i.e., birds that winter in warmer climates than their breeding range (typical of most migratory birds; Dufour et al., 2020), can be distinguished from temperate sedentary birds, that, oppositely, generally experience warmer conditions during growth. However, niche-switching migratory birds cannot be reliably distinguished from the tropical sedentary birds born near the wet season. This observation also suggests reduced reliability of the test in monsoon-driven regions (e.g., East or South Asia), where sedentary individuals generally hatch during the rainy season (Sundar et al., 2018). Niche-tracking migratory birds, i.e., those following consistent climatic conditions year-round, cannot be differentiated from sedentary birds living in regions with low climate seasonality. Lastly, migratory birds wintering in colder climates (e.g., short-distance, longitudinal, or possibly some altitudinal migrants, not illustrated here) are indistinguishable from sedentary birds living in temperate climates or experiencing cooler non-breeding temperatures in subtropical or tropical regions.

##### (ii) Non-Local Origin Test

At the site level, a bird can be identified as non-locally born if its δ^18^O_p, EBT_ value differs significantly from those of locally born individuals (juveniles or sedentary). This enables the identification of migratory birds that were born in a different (typically colder) climate, and died during wintering or at migratory stopovers.

Below, we discuss the biological and environmental sources of variability in δ^18^O_p_ and potential confounding factors likely to affect the sensitivity (i.e., detectability of migratory behaviours) and specificity (i.e., reliability of inferences) of these two tests, and outline how appropriate sampling strategies can help mitigate this variance and maximise classification accuracy.

#### 4.1.2 On bone remodelling dynamics

Among other processes, cortical bone remodelling – responsible for LBT deposition – occurs as a reparative response to microcracks caused by biomechanical stress (Lee et al., 2002; Ponton et al., 2007; Mitchell et al., 2017), or as a result of bone resorption triggered by high metabolic demands, such as those following molting in some species (Meister, 1951). These processes are sensitive to life history traits and environmental factors, raising the possibility of seasonally biased LBT deposition. Whether such biases vary across individuals or species with different physiologies, phenologies, or locomotor behaviours (e.g., migratory vs. sedentary) remains largely unexplored.

Our simulation framework showed that if LBT were deposited exclusively right after breeding – e.g., during a single molt phase post-breeding – this would represent a worst-case scenario, minimising isotopic differences between EBT and LBT and dramatically reducing the detectability of migratory behaviour using the *Climatic Niche Contrasts Test*. However, empirical data we obtained for both sedentary and migratory birds sometimes showed large contrasts in δ^18^O_p_ values between EBT-rich and LBT-rich samples, inconsistent with a strictly post-breeding deposition of LBT across species. Overall, extreme temporal biases in LBT deposition are unlikely given the diversity of underlying factors described above. This is even less likely for migratory species, which frequently exhibit stepwise molt patterns – initiating the molt cycle on breeding grounds and completing it on wintering grounds (Kjellén, 1994; Zuberogoitia et al., 2018) – and for which more frequent remodelling events can be expected around bio-mechanically and metabolically costly migratory journeys. Thus, in both sedentary and migratory species, LBT deposition is likely more evenly distributed over the lifespan, which should yield empirical resolution for the *Climatic Niche Contrasts Test* closer to that estimated under the assumption of random deposition.

Additionally, bone remodelling rates are known to depend on body size and metabolic demand (de Buffrénil & Quilhac, 2021b), which could explain why the smaller species in our sample (*C. corone* and *F. subbuteo*) tended to exhibit a lower proportion of LBT. This may reduce the likelihood of capturing LBT deposited outside the breeding season – if such events did occur – and therefore lower the sensitivity of the *Climatic Niche Contrasts Test* for smaller birds. Increasing sampling effort for such species may help mitigate this limitation.

#### 4.1.3 Non-migratory movements: potential confounding factors?

In both sedentary and migratory species, there exist a range of non-migratory movements that differ from regular seasonal migrations between breeding and non-breeding grounds. These include foraging movements, nomadism (i.e., population-level shifts triggered by unpredictable resource scarcity), and dispersal movements – either from natal site (natal dispersal) or from a breeding site to another (breeding dispersal) (Dufour et al., 2023). Among these, natal dispersal appears as the most likely confounding factor for migration inference, as it typically occurs shortly after EBT deposition and is widespread among vertebrates. However, our simulations showed that climatic conditions remained relatively similar before and after natal dispersal, especially across typical dispersal distances (< 20 km; Paradis et al., 1998; Claramunt, 2021; Fandos et al., 2023) and even under extreme scenarios (several hundred kilometres; Solheim & Stefansson, 2016; up to 1000 km), resulting in minimal impact on the *Climatic Niche Contrasts Test*.

More exceptionally, vagrancy – defined as movements beyond a species’ typical range – can result in larger-scale non-migratory movements, sometimes over thousands of kilometres, especially in lighter bird species more susceptible to wind-drift or rafting (Dufour et al., 2023). Although rare and typically less pronounced in sedentary species than in migratory ones, vagrancy could theoretically lead to false positives (i.e., misclassification of sedentary birds as migratory) under both the *Climatic Niche Contrast Test* and the *Non-local Origin Test*, particularly if it occurs early in life, prior the deposition of most LBT patches. However, such misclassifications could be detected through careful observation and mitigated by appropriate sampling strategies. First, vagrants usually occur only sporadically at a site. Second, if results across several individuals from the same population are inconsistent – reflecting diverse and directionally variable origins – this would likely suggest recurrent vagrancy rather than structured migratory behaviour. For this reason, applying the *Climatic Niche Contrast Test* at the population level could help average out rare stochastic movements (i.e., randomly distributed within populations) and maintain the test’s specificity for detecting niche-switching migration.

#### 4.1.4 Population-level inference: averaging out random effects

Our simulations highlighted that applying the *Climate Niche Contrast Test* at the population level (i.e., using samples from multiple conspecifics at the same locality) can substantially improve the reliability of inferences. This is because many sources of variation in life histories or climate (among others) and resulting δ^18^O_p_ values are stochastic rather than systematic. Population-level sampling therefore mitigates the influence of individual “outliers”, as suggested for vagrancy in 4.1.3. Other notable examples include temperate sedentary birds that hatched either during exceptionally cold years or unusually early in the breeding season – while temperatures were still relatively low (e.g., as early as February-March in some species) – and that could otherwise be misclassified as migratory if assessed individually.

Another source of stochasticity stems from physiological and ecological processes affecting isotopic discrimination between δ^18^O in precipitation (δ^18^O_w_) and in bone phosphate (δ^18^O_p_). As Wunder and Norris (2019) aptly noted, “birds are not rain gauges”. Factors such as age, sex, metabolic rate, foraging behaviour, mixed water sources, water reservoir turnover rates, or enrichment of heavy isotopes due to evaporation can all influence tissue δ^18^O_p_ values (Koch, 1998; Bowen & West, 2019; Hu et al., 2023). In our framework, we assumed these effects remain stable during the formation of a given tissue (e.g., EBT or a single bone remodelling event) but vary randomly across an individual’s lifetime. While such residual variation can locally weaken or even reverse the otherwise positive correlation between δ^18^O_w_ and δ^18^O_p_ in some individuals, these deviations tend to cancel out when averaged across a population, thereby further reducing the risk of misclassifying sedentary birds as migratory (i.e., false positives). However, if the entire population is uniformly exposed to the same bias (e.g., all individuals seasonally relying on the same evaporated water source), residuals may not cancel out, reducing the specificity of the population-level *Climate Niche Contrast Test* compared to our simulation expectations. Nevertheless, because the various contributing processes are unlikely to systematically bias all individuals in the same direction, population-level application is still expected to yield more reliable inferences than individual-level analyses.

#### 4.1.5 Experimental validation

We here applied both the *Climatic Niche Contrast Test* and the *Non-local Origin Test* to a small set of extant adult birds, comprising four migratory and three sedentary individuals. Two migratory individuals had died outside the documented breeding ranges of their species, but only one showed sufficient isotopic contrast between hatching and death locations during the growth season to be confidently inferred as non-local (*G.grus*). Among the four migratory birds, only one was confidently identified as migratory by the *Climatic Niche Contrast Test (G.grus)*, and, importantly, no sedentary individuals were misclassified as migratory. The limited detection of some migratory individuals using the *Climatic Niche Contrast Test* could be explained by the poor climatic contrast between their breeding and wintering grounds. Indeed, the death locations of these individuals suggest they belonged to populations migrating over relatively short latitudinal ranges – likely no more than 15° to 20° – which may be insufficient to produce a strong isotopic contrast. Alternatively, the absence of migratory signal could simply reflect an “unlucky” sample that, by chance, did not include LBT deposited during migration or wintering.

While these empirical applications provide encouraging qualitative support for the approach, they do not constitute formal experimental validation, which is, in practice, hardly attainable given current material and logistical constraints. Robust empirical validation would require: (i) sample sizes that are difficult to obtain from comparative anatomy collections, where destructive sampling is discouraged for species with few available specimens; (ii) access to multiple individuals per species from the same locality; and, ideally, (iii) detailed life history data for each sampled individual, including migratory routes and experienced climatic conditions. In the future, this latter point could, for instance, be addressed through opportunistic sampling of birds that died during or after tracking. Even so, at the time of this study, both the cost and time demands of the protocol remain a considerable challenge for such large-scale implementation.

Given these limitations, an empirically-grounded simulation framework remains the most feasible and robust approach for assessing the performance of the two tests introduced in this study. Across all modelled scenarios – spanning both optimistic and pessimistic assumptions – false positive remained negligible under the recommended sampling strategies (4.1.1 - 4.1.4). We therefore argue that, under these conditions – including an applicability currently restricted to temperate climates for the *Climatic Niche Contrast Test* – both tests can already be applied empirically with minimal risk of mistaking sedentary birds for migratory. In the future, these models, and hence the expected performances of the proposed tests, could: (i) be refined as new physiological insights emerge, particularly regarding LBT deposition dynamics (4.1.2) and the metabolization of rainwater within bone tissues of wild bird species (4.1.4); (ii) be further evaluated under extant climates not represented in this study (e.g., monsoon-driven regions); and (iii) even integrate models of past climate seasonality relevant to any geological time of interest (see 4.2).

### 4.2 Transposing this approach to deep time

The tests outlined in Section 4.1 provide a framework for identifying migratory behaviours in extant bird species through δ^18^O_p_ analysis of bones tissues. However, their application to fossil specimens introduces several critical considerations.

#### 4.2.1 Diagenetic alteration

Fossil bones can be subject to diagenetic alteration, particularly through interactions with meteoric waters during fossilisation, which can modify the original δ^18^O_p_ values and potentially undermine the reliability of migratory behaviour predictions. Nonetheless, phosphate apatite has been shown to routinely preserve pristine oxygen isotope compositions in fossil bones and teeth dating back to the Mesozoic (Amiot et al., 2011; Angst et al., 2014; Amiot et al., 2017), when birds arose among theropod dinosaurs. Well-established diagnostic methods (Iacumin et al., 1996; Lécuyer et al., 2003; Pucéat et al., 2004; Zazzo et al., 2004; Kral et al., 2024) can effectively assess and mitigate the impact of such alterations, preserving the integrity of isotopic analyses. Notably, diagenetic alteration can be monitored by measuring the δ^18^O values of both phosphate and carbonate fractions of bone apatite, which are expected to be highly correlated in non-altered bones (Iacumin et al., 1996). Furthermore, numerous studies have demonstrated that bone microstructure can be exceptionally well preserved through fossilisation (Chinsamy-Turan, 2005; de Buffrénil et al., 2021), supporting the feasibility of measuring EBT and LBT proportions in fossil specimens.

#### 4.2.2 Sampling constraints

The histological analyses required for the *Climatic Niche Contrast Test* can only be reliably conducted on well-preserved three-dimensional bones, which are rarely found in anatomical connexion at fossil sites. Even when found in close proximity, fossil specimens assigned to the same species are typically assumed to represent different individuals. Since the sampling of multiple bones is necessary to achieve sufficient sample sizes and access both EBT-rich and LBT-rich samples – at least given the precision of the sampling techniques employed in this study – this restricts the *Climatic Niche Contrast Test* to population-level applications. While population-level sampling for extant specimens can be limited by the scarcity of individuals from the same locality in comparative anatomy collections, as was the case in this study, this is not a significant limitation at many fossil sites, where abundant material from certain species is often available (Olson, 1975; Cheneval, 1989; Mourer-Chauviré, 2006; Mayr, 2020; Núñez-Lahuerta et al., 2024).

The *Non-local Origin Test* (Section 4.2) relies on accurate controls for local breeding season δ^18^O_w_ values. For fossil sites, these controls can ideally derive from juvenile bones – readily identifiable by their shape and surface texture (Watanabe, 2018) – or from EBT-rich samples of bird species known to be sedentary (e.g., flightless taxa). In some well documented fossil sites, such material is abundant and spans a wide variety of bird species (e.g., Olson, 1975; Cheneval, 1989). In the absence of sedentary or juvenile birds, alternative proxies for growth season temperatures, such as the δ^18^O of fossil plants or fossils of sedentary and/or juvenile vertebrates (e.g., mammal teeth; Bernard et al., 2009), could theoretically serve as substitutes. However, these alternative proxies would require careful calibration to ensure they reflect geographic and temporal signatures comparable to those recorded in bird bone tissues (Section 4.1.2). Incorporating these proxies would likely introduce additional variability to the local growth season δ^18^O_w_ values, thereby reducing the resolution of the test.

#### 4.2.3 Evolving δ^18^O_w_ isoscapes

As precipitation δ^18^O_w_ isoscapes evolve over geological time in response to changing climatic conditions (Bowen, 2010), prior knowledge of the paleoclimatic and stratigraphic context of the fossil site is required to ensure the efficacy of the tests. The *Climatic Niche Contrast Test* is primarily suited for fossil sites associated with temperate climates, because although it effectively differentiates niche-switching migratory populations from temperate sedentary populations, it cannot distinguish them from tropical sedentary ones with confidence (Section 4.1). Applying this test to tropical contexts – or, similarly, to monsoon-driven localities (not explored here, 4.1) – would therefore risk significant misclassification of sedentary populations as migratory. Furthermore, both tests – *Climatic Niche Contrast Test* included when performed at the population level – are likely to decrease in resolution if applied to a fossil site exhibiting high stratigraphic uncertainty, especially if it spanned significant climatic changes over its stratigraphic range. Similarly, the resolution of both tests would inevitably decrease if applied to periods with lower global latitudinal δ^18^O_w_ gradients and/or reduced climate seasonality, such as the Miocene Climate Optimum (Herold et al., 2011). These concerns can be mitigated by integrating stratigraphic and paleoclimatic data from existing literature for the relevant geological periods and sites.

### 4.3 Conclusion and future directions

Overall, combining histological analyses with oxygen isotope analyses of bone tissues appears to provide a reliable tool for predicting migratory behaviours in extant birds and holds significant promise for fossil applications, provided that the palaeoclimatic context is carefully accounted for. For now, this approach provides a predictive power limited to specific migratory behaviours (i.e. niche-switching and/or migratory birds that died outside their natal sites), and an applicability sometimes limited to specific climatic contexts (e.g. temperate climates). Sensitivity could be further improved by adopting a multi-proxy isotopic framework (Wunder & Norris, 2019; Hobson & Kardynal, 2023), while maintaining awareness of the spatio-temporal context of bird bone mineralisation emphasised in this study. In particular, combining a meteoric water-based proxy (δ^18^O or δ^2^H) with another complementary spatial proxy – such as plant-based (δ^13^C) or bedrock-based (^87^Sr/^86^Sr) proxies – could enable the identification of migratory behaviours that are not or poorly detected by a δ^18^O analysis alone (e.g., longitudinal or niche-tracking migrations).

Applied to fossil specimens from sites spanning the Cenozoic, or even as far back as the Late Cretaceous (where bird nesting traces have been reported in polar ecosystems; Wilson et al., 2025), this approach could be key to unravelling the origin(s) and evolution(s) of migratory behaviour in birds, particularly by providing spatiotemporal snapshots of past migratory behaviours that would enable the calibration of methods of ancestral state or niche reconstruction across the phylogeny of birds. More broadly, this study establishes a framework for inferring migratory behaviour in any vertebrate, based on non-fully remodelling biomineralised remains, whether extant or fossilised.

## Author contributions

AD, AL, RA, AVL designed the project. AD, RA, AVL and FF acquired the data. AD, RA, and AC analysed the experimental data. AD and JJ designed the theoretical model and statistical analyses. AD wrote the first draft. All authors contributed to revisions.

## Acknowledgements

This research received funding from the CNRS-INSU TelluS-INTERRVIE programme in 2022 (“IsoMig” project) and 2023 (“E-VOL” project). The authors would like to thank the following people for their valuable contributions, despite the difficulties caused by the Covid crisis: Emmanuel Robert and Didier Berthet for providing access to the extant and/or fossil specimens from the Université de Lyon 1 and the Musée des Confluences respectively; Colette Guilbaud, Christophe Névado and Doriane Delmas for the preparation of thin sections; Magali Séris for completing the preparation of some chemically treated samples; Vivian de Buffrénil and Jessica Mitchell for providing advice on bird bone histology.

## Data availability

All data necessary for the reproducibility of this study—including R scripts, theoretical and experimental datasets—are available in the following Zenodo repository: https://doi.org/10.5281/zenodo.14825377

## Conflict of interests

The authors declare no conflict of interest.

## Statement of inclusion

In this study, we focus primarily on the African-European geographic domain, both for skeletal specimens access (experimental approach) and computational time (theoretical approach) convenience. Future research could extend this work by incorporating data from other regions, further enhancing the global perspective we seek to achieve.

## SUPPORTING INFORMATIONS OF

**FIGURE S1:**
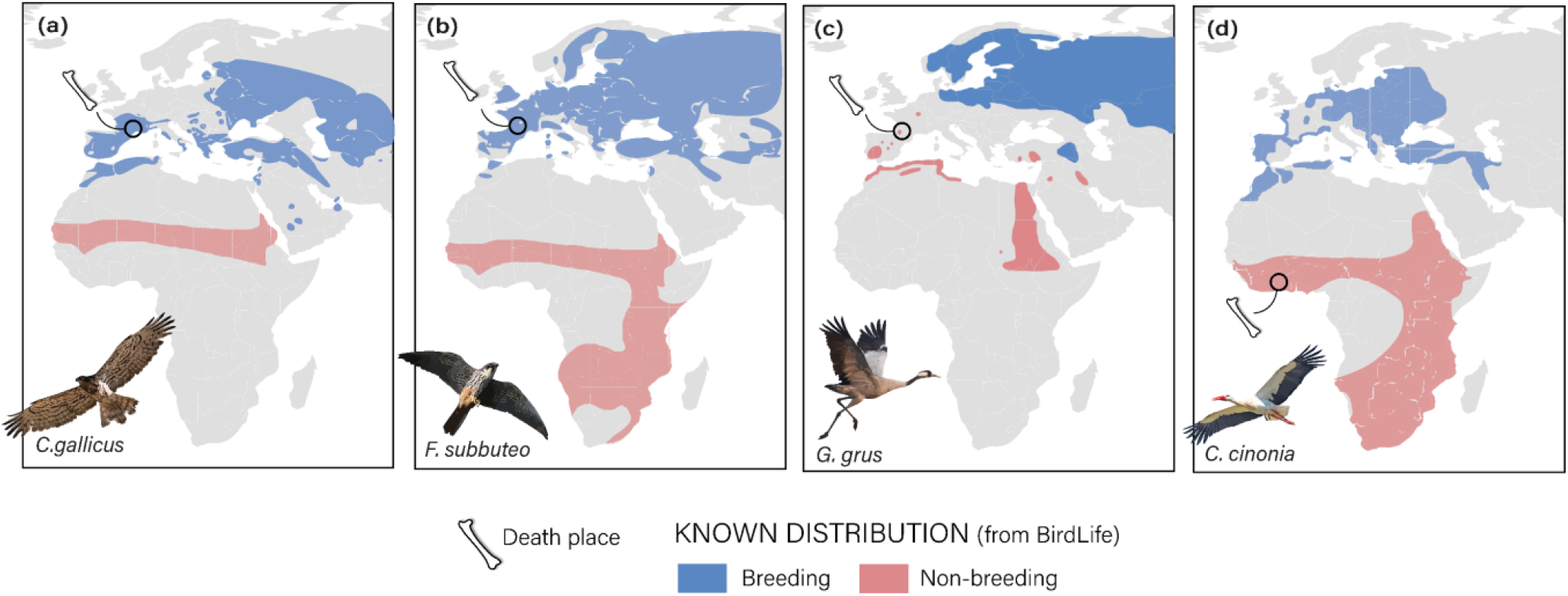
Death locations of migratory individuals from the application case, and known breeding and non-breeding areas of their species (modified after eBird data maps; Sullivan et al., 2009).

## APPENDIX A.

Preliminary tests protocols to control for potential biases on bones isotopic composition

### A.1 Test the influence of sodium perborate treatment on bone isotopic value

A sodium perborate treatment was historically performed to clean the skeletal specimens of the Collections de l’Université de Lyon 1 from their tendinous and cartilaginous parts. We therefore controlled the influence of these chemical – which contains oxygen – on the oxygen isotope ratios of bone tissues. We compared the oxygen isotope ratio of two subsamples deriving from the same bone sample of a Great cormorant: one of it treated using sodium perborate, the other remaining untreated. The sodium perborate treatment consisted in boiling the bone sample several hours in a 2 g.L^−1^ boiling sodium perborate solution. This treatment appears to have no significant effect on the oxygen isotope discrimination of a test individual’s bones (Fig S1).

### A.2 Skeleton isotopic cartography to test potential heterothermies

Oxygen isotopic fractionation is known to be temperature-dependent. Given that birds are endotherms, and that some bones are surrounded by more soft tissues than others, fleshy body-parts are supposed to be controlled by the constant body temperature (around 40°C in birds), while thinner body parts may approach ambient air temperature. We therefore carried out an isotopic mapping of the skeleton of a juvenile bird to test whether potential heterothermies at the skeletal scale affect the oxygen isotopic fractionation during bone mineralisation. The control specimen was chosen juvenile to avoid biases associated with bone remodeling, which can vary between bones. This test was run on 6 different post-cranial bones of a juvenile extant chicken (Table S1, Fig S1, A). As we found no significant difference in oxygen isotope fractionation between the bones from more or less fleshy body parts (Fig S1), isotopic fractionation was considered homogeneous at the individual level, regardless of potential heterothermies within the body.

**FIGURE S2:**
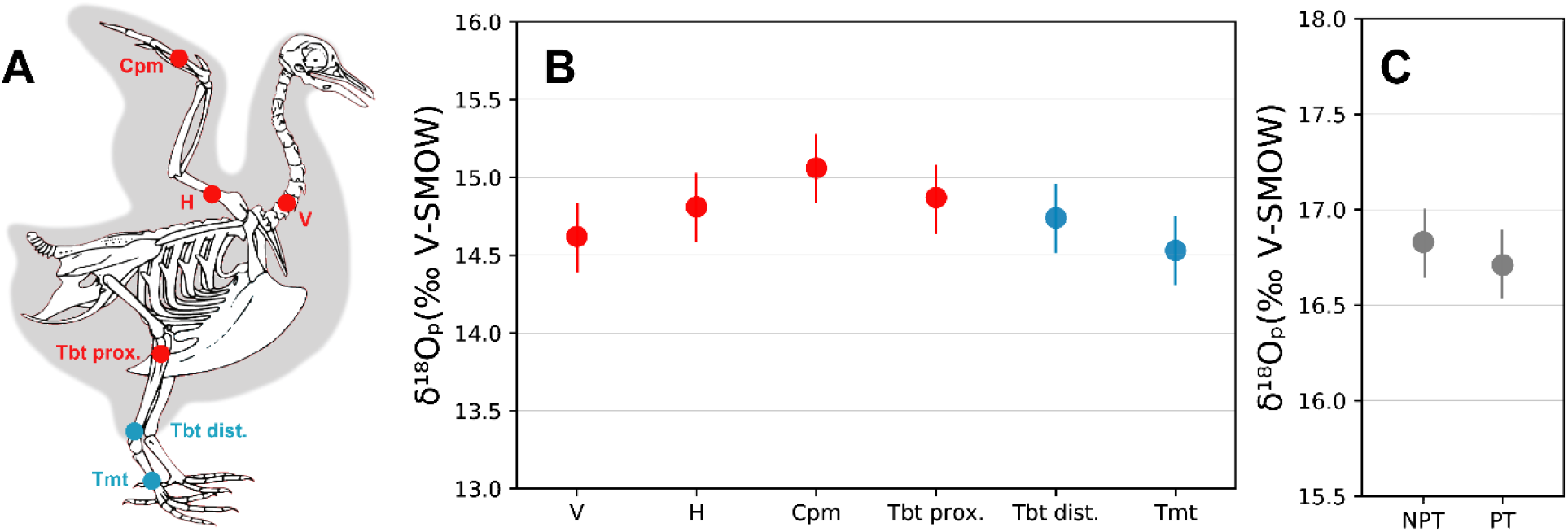
Influence of heterothermy (B) and sodium perborate treatment (C) on bones’ δ^18^O_p_. A: shows the fleshiness of the chicken body parts surrounding the bone tested for the heterothermy test, red points for fleshy parts and blue point for relatively less fleshy parts on both A and B. Abbreviations: V, vertebrae; H, humerus; Cpm, carpometacarpus; Tbt prox., proximal tibiotarsus; Tbt dist., distal tibiotarsus; Tmt, tarsometatarsus; NPT, no perborate treatment; PT, perborate treatment. Error bars are the mean analytical error for all chemistries (±0.22 ‰).

## APPENDIX B.

Diagnostic criteria to identify and quantify histological features on thin sections

Secondary osteons can be distinguished from primary osteons based on the several characteristics, as follows:

i. They are surrounded by a cement line (Fig. S2, b, d), which is the result of their particular formation process. Indeed, the cement line results from the local resorption of bone by osteoclasts in the cortex, and the subsequent centripetal deposition of secondary bone by osteoblast cells. The cement line is scalloped, delimitates each individual secondary osteon, and marks the interface between tissues deposited at different time intervals. By definition, primary osteons are not surrounded by a cement line.
ii. This continuous process of resorption-redeposition, characteristic of bone remodeling, can also have for consequence a possible overlap between different secondary osteons generations as shown in Fig. S2, f, and an overlap on underlying primary bone tissues.
iii. Moreover, remodeling and the associated formation of secondary osteons are usually initiated in the peri-medullary, deep cortical region and extend progressively towards the periosteal surface throughout the life of an individual. In birds, secondary osteons are thus preferentially concentrated in the deep and mid cortex, whereas primary osteons found in the primary woven-parallel complex form more homogeneously throughout the cortex.
iv. Another characteristic feature is that in birds, in general, secondary osteons are larger in diameter than primary osteons and also that their shape is less homogeneous.
v. Finally, secondary osteons are formed of a parallel-fibered or lamellar bone matrix and thus have a different texture than the surrounding woven bone under both natural and polarized light (Fig. S2, a, c, e, f).

Note that Maltese crosses, which are artefacts visible on osteons in polarized light (Fig. S2, e), are often considered as a diagnostic criterion for secondary osteons, but were here occasionally observed on primary osteons.

Most of the primary periosteal cortex is formed of a well-vascularized woven-parallel complex (woven bone matrix associated to primary osteons) in birds. During growth, there is also centripetal deposition of an internal fundamental system (IFS) that marks the end of the medullary cavity expansion. This layer is made of an endosteal (often avascular) lamellar bone tissue and is secondary in nature. Towards the end of growth, there is formation of an external fundamental system (EFS) that marks the attainment of skeletal maturity. The EFS is composed of poorly vascularized to avascular primary parallel-fibered bone in most cases. The EFS and IFS are easily distinguishable from other bone features (such as secondary osteons) and appear clearly birefringent under polarized light (linked to the orientation of collagen fibers and hydroxyapatite crystals, Chinsamy, 2005, Fig. S2, c, e, f). In rare cases, secondary osteons can form before the end of the chick’s development (so before the EFS formation) to repair fractures, micro-damages, or because the chick went through an unusual stress-inducing event. This early remodeling event can be recognized by secondary osteons being cross-cut by the IFS (Fig. S2, f), but is hard to quantify because of the possible overlying of several generations of secondary osteons. It is therefore a potential source of error on proportion determination.

**FIGURE S3:**
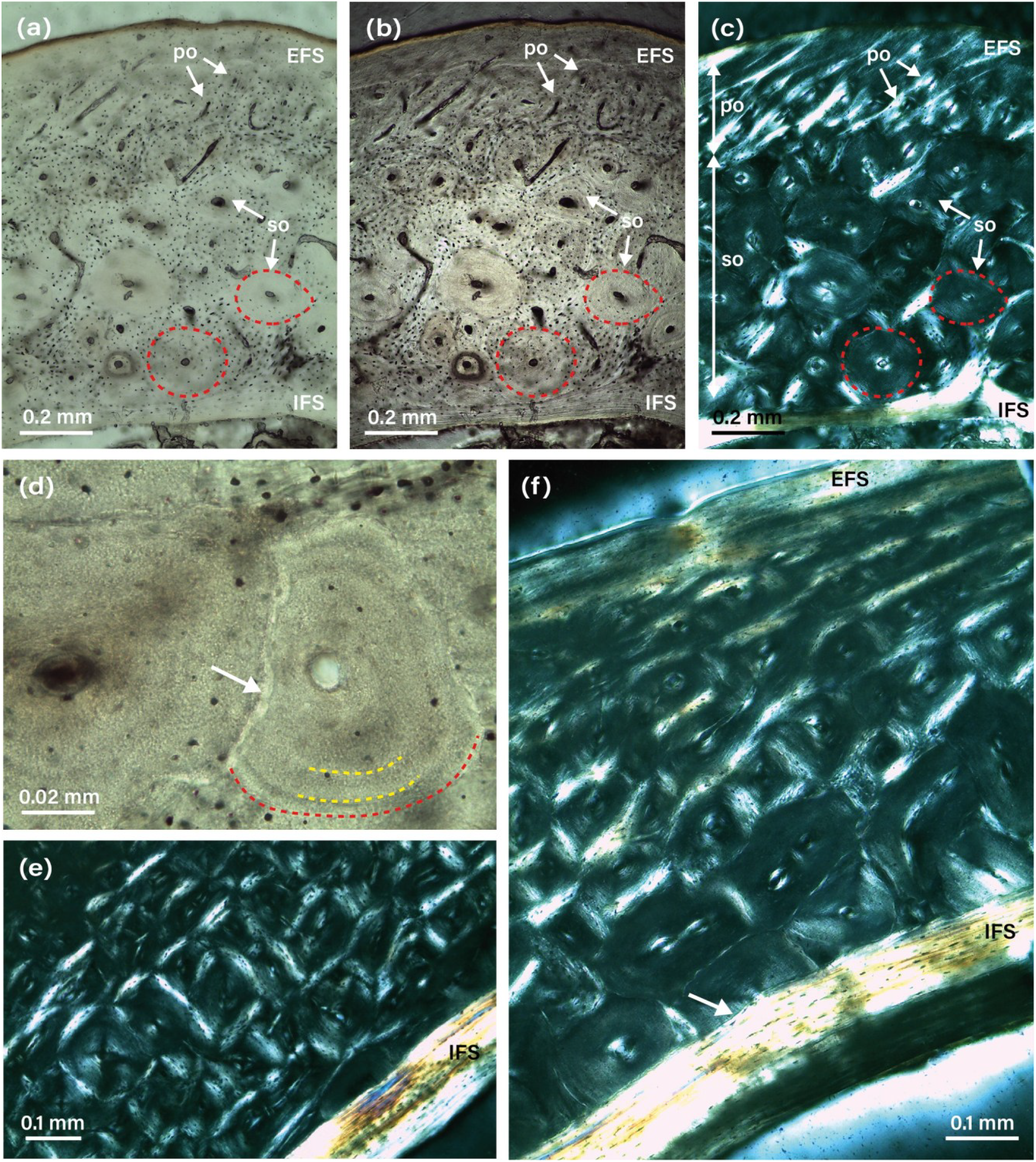
Natural (a, b, d) and polarized light (c, e, f) images showing bone tissues in extant adult birds, illustrating the diagnostic criteria developed in Appendix A. (a), (b) and (c) show the same region of a White Stork (*Ciconia ciconia*) tibiotarsus under different lights, and a zoom on secondary osteons. (d) shows the cement line delimiting the osteons (red dotted line), lamellae (yellow dotted lines), and an overlapping relationship between osteons (white arrow). Note that limiting the transmitted light amount with the microscope condenser makes secondary osteons cement line more visible (b, d), and that primary fibrolamellar bone between primary (po) and secondary osteons (so) appears highly birefringent in polarizing light. Internal (IFS) and external fundamental systems (EFS) appear also very bright in polarizing light. (e) *Grus grus* tibiotarsus detail showing maltese crosses within secondary osteons. (f) *Grus grus* tibiotarsus showing secondary osteons cut by the IFS, evidence that this bone remodelling took place when the bird was juvenile. Conventionally, all the images are oriented with the medullary cavity on the bottom.

**FIGURE S4:**
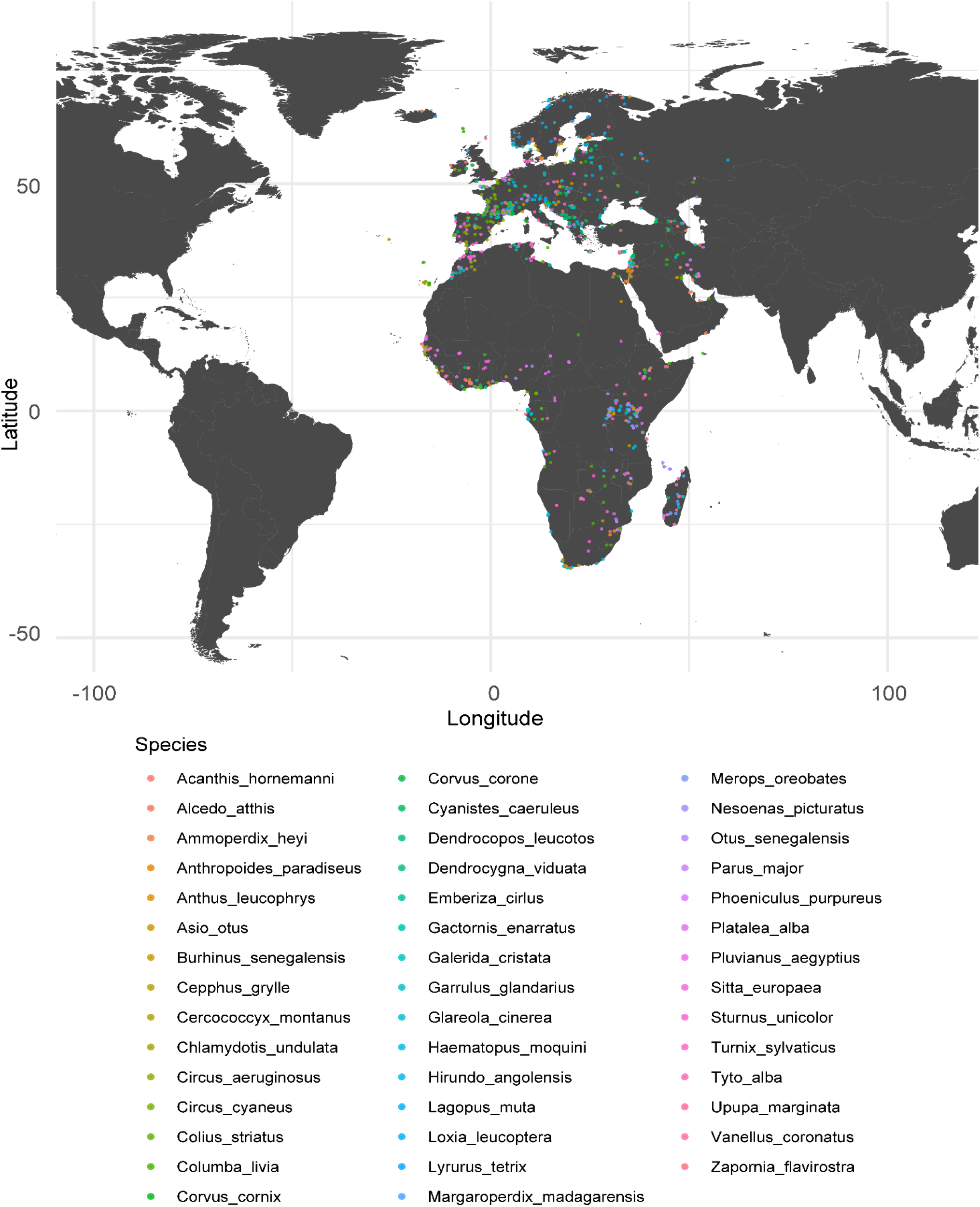
Map depicting year-round living location of sedentary individuals, drawn from the revised 2023 eBird Basic Dataset. Individuals are color-coded by species.

**FIGURE S5:**
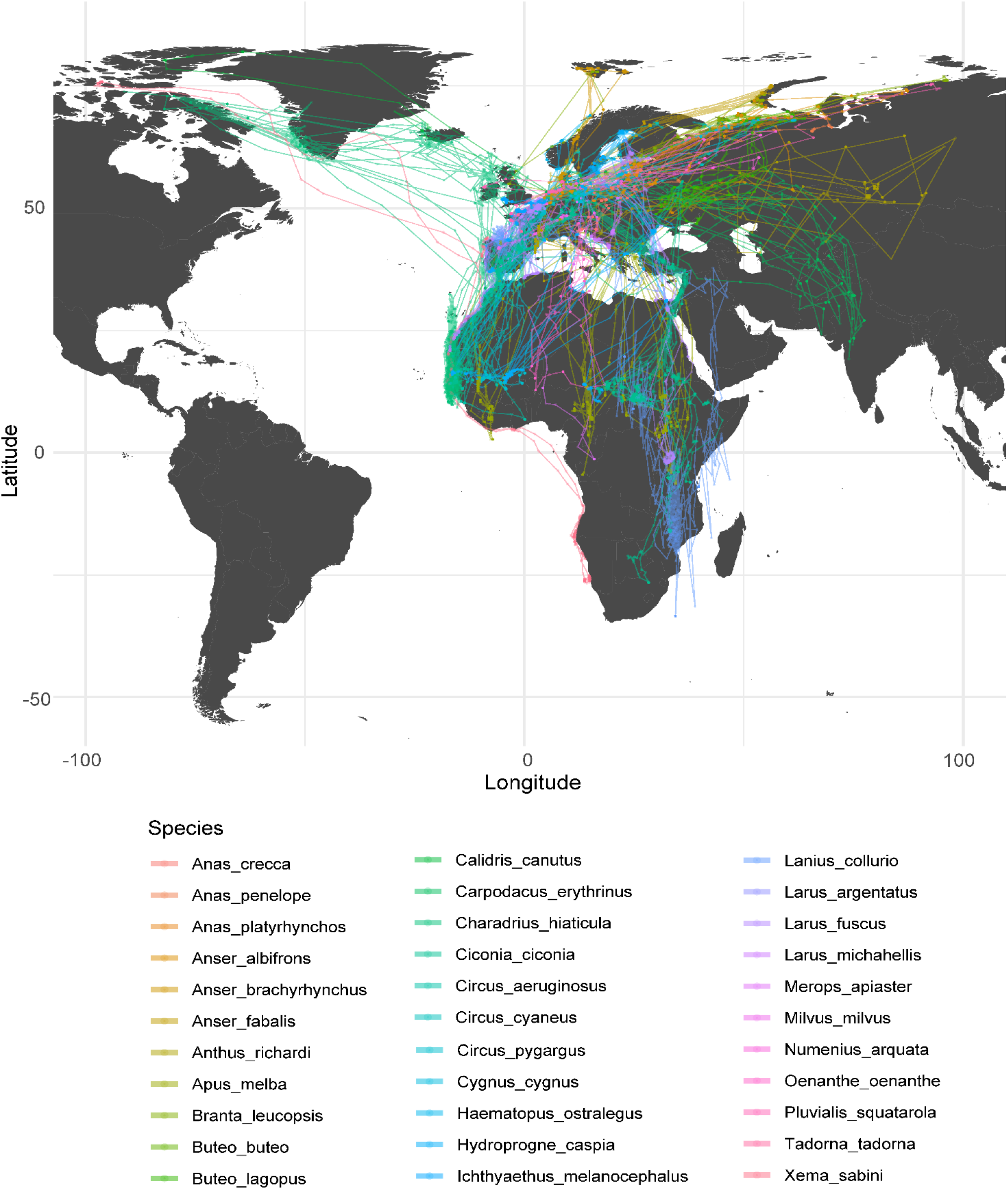
Map depicting one year of migratory routes for each individual, derived from data available in the Movebank.org repository (see Data Source section and Table S1). Individuals are color-coded by species.

## APPENDIX C.

Fractionation equation under comparison

We fitted a RMA regression (Frac. Eq. 1) between the δ^18^O_p_ measured on juvenile chicken bones and the mean annual δ^18^O_w_ values in precipitations extracted from the Waterisotope portal (http://waterisotopes.org, Bowen & Revenaugh, 2003). However, hatching dates available in Amiot et al. (2017) indicated that most of these juvenile specimens lived during the season of lowest local δ^18^O_w_ values in precipitations (Table S1), making annual mean δ^18^O_w_ value in precipitations a potentially biased (overestimated) estimate of the bird drinking water δ^18^O_w._

We therefore revised Amiot et al. (2017) oxygen isotope fractionation equation by re-estimating juvenile chickens’ drinking water δ^18^O_w_ values as the local δ^18^O_w_ in precipitations averaged over the 30 days post-hatching. We first fitted a RMA regression model on the revised data (Frac. Eq. 2). Then, to ensure better calibration against wild birds from various species, we fitted a RMA regression model on the revised data incorporating to the dataset the measured δ^18^O_p, EBT_ values of juvenile and sedentary birds, and modelling their mean δ^18^O_w, EBT_ values as in section 2.2.2.1 (Frac. Eq. 3). Finally, we tested a last equation by keeping the δ^18^O_w_ of chickens published by Amiot et al. (2017), but including juvenile and sedentary bird samples from this study with δ^18^O_w, EBT_ values modelled as in section 2.2.2.1 (Frac. Eq. 4).

We simulated all plausible δ^18^O_p,EBT_ values for the juvenile and/or sedentary birds sampled (2.1.1), assuming their death sites matched their hatching sites, and all plausible δ^18^O_p,LBT_ values for the somatically mature sedentary birds sampled (2.1.1), assuming their death sites represented year-round living locations. These values were obtained following the methodology described in 2.2.4. No δ^18^O_p_ values were modelled for the somatically mature migratory birds sampled due to unknown birthplaces and exact migratory routes. We excluded δ^18^O_p,LBT_ computations for samples with less than 20% LBT due to high cumulated errors in equation 1. Because all fractionation equation gave similar distribution, we kept equation Frac. eq. 3 for the remainder of the manuscript.

Please note that the δ^18^O_p_ measurements used in the present paper to extend the dataset of Amiot et al. (2017) were obtained using the same reference materials as in that study (see Main Text, section 2.1.4). If reference materials with different assigned values are employed to further refine or revisit this equation, it becomes necessary to recalibrate the measurements using the values described in Section 2.1.4 (Main Text) to ensure data comparability. Likewise, calibration to the values reported in Section 2.1.4 is required if one aims to (i) estimate δ^18^O_w_ from δ^18^O_p_ measurements or (ii) directly compare δ^18^O_p_ values with those reported in our study. However, if future users aim solely to predict migratory status following our experimental framework (Section 2.1, Main Text) – that is, by applying tests directly to δ^18^O_p_ measurements without comparison to other datasets – there is no need to re-calibrate measurements to align with our reference materials.

**FIGURE S6:**
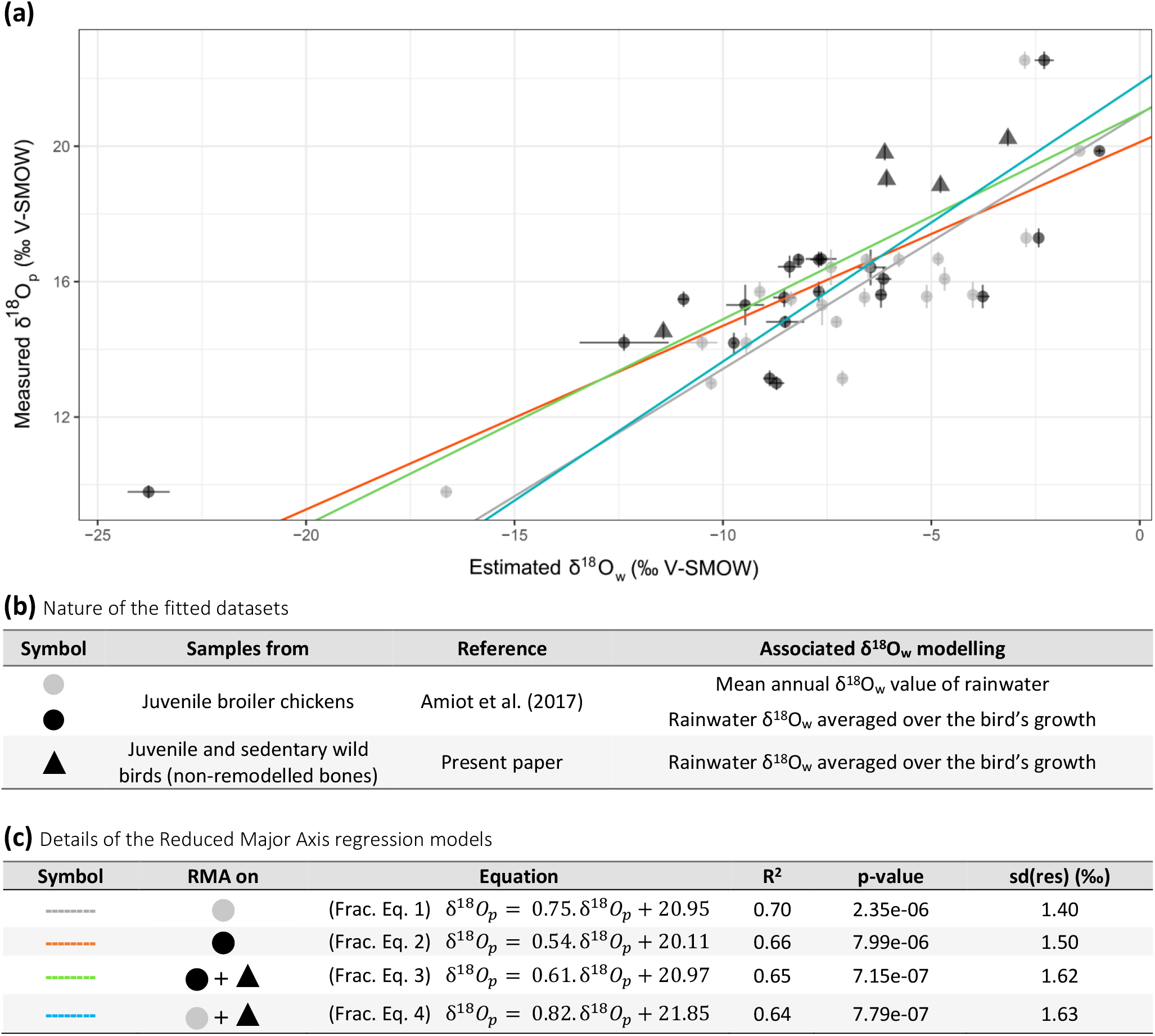
Comparison of fractionation equations linking drinking water δ^18^O_w_ estimates to bone phosphate δ^18^O_p_ measurements in birds, using reduced major axis (RMA) regression. The fit was performed either on the raw δ^18^O_p_ and δ^18^O_w_ data from Amiot et al. (2017), on the δ^18^O_p_ values from Amiot et al. (2017) against revised δ^18^O_w_ estimates, or on the same dataset extended with control data from the present study (juvenile and sedentary birds). The nature of the fitted datasets is described in panel (b), and details of the regression models are provided in panel (c).

**FIGURE S7:**
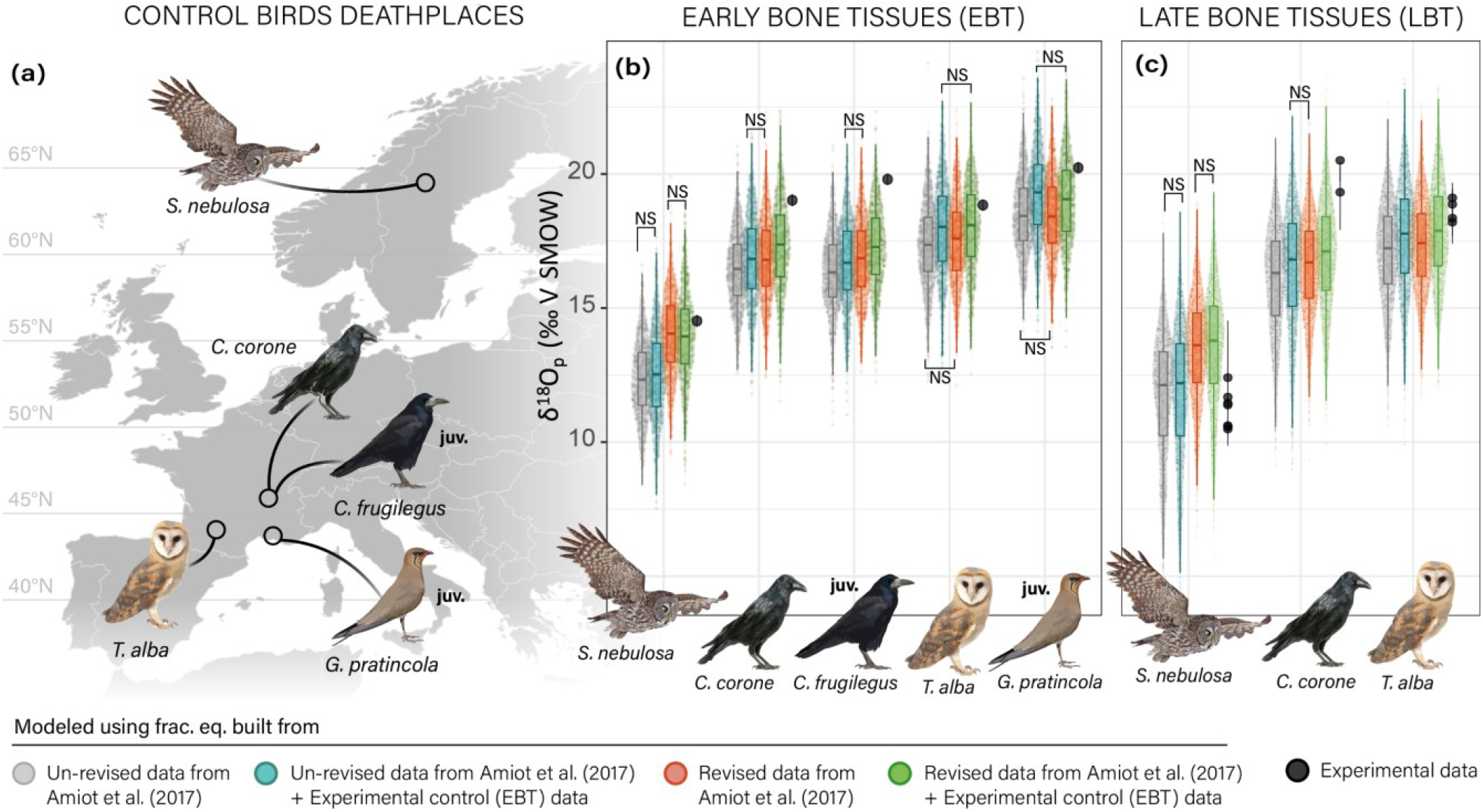
For each sedentary and/or juvenile bird sampled: (a), death location; for EBT (b) and LBT (c), comparison of experimental tissue-specific δ^18^O_p_ values with values modelled using either unrevised or revised fractionation equations (see Fig. S6). Each modelled δ^18^O_p,EBT_ value corresponds to a 30-day growth window, while each modelled δ^18^O_p,LBT_ value represents a 10-day bone remodelling event. Experimental δ^18^O_p,EBT_ values are derived from a single bone sample and its corresponding thin section. Error bars on experimental δ^18^O_p,EBT_ values (smaller than dots’ sizes) show mean analytical error for mid-ulnae δ^18^O_p,bulk_ measurements. Error bars on δ^18^O_p,LBT_ values reflect propagated mean analytical error from δ^18^O_p,bulk_ to δ^18^O_p,LBT_ via equation 1. Juvenile specimens are labelled “juv.”.

**FIGURE S8:**
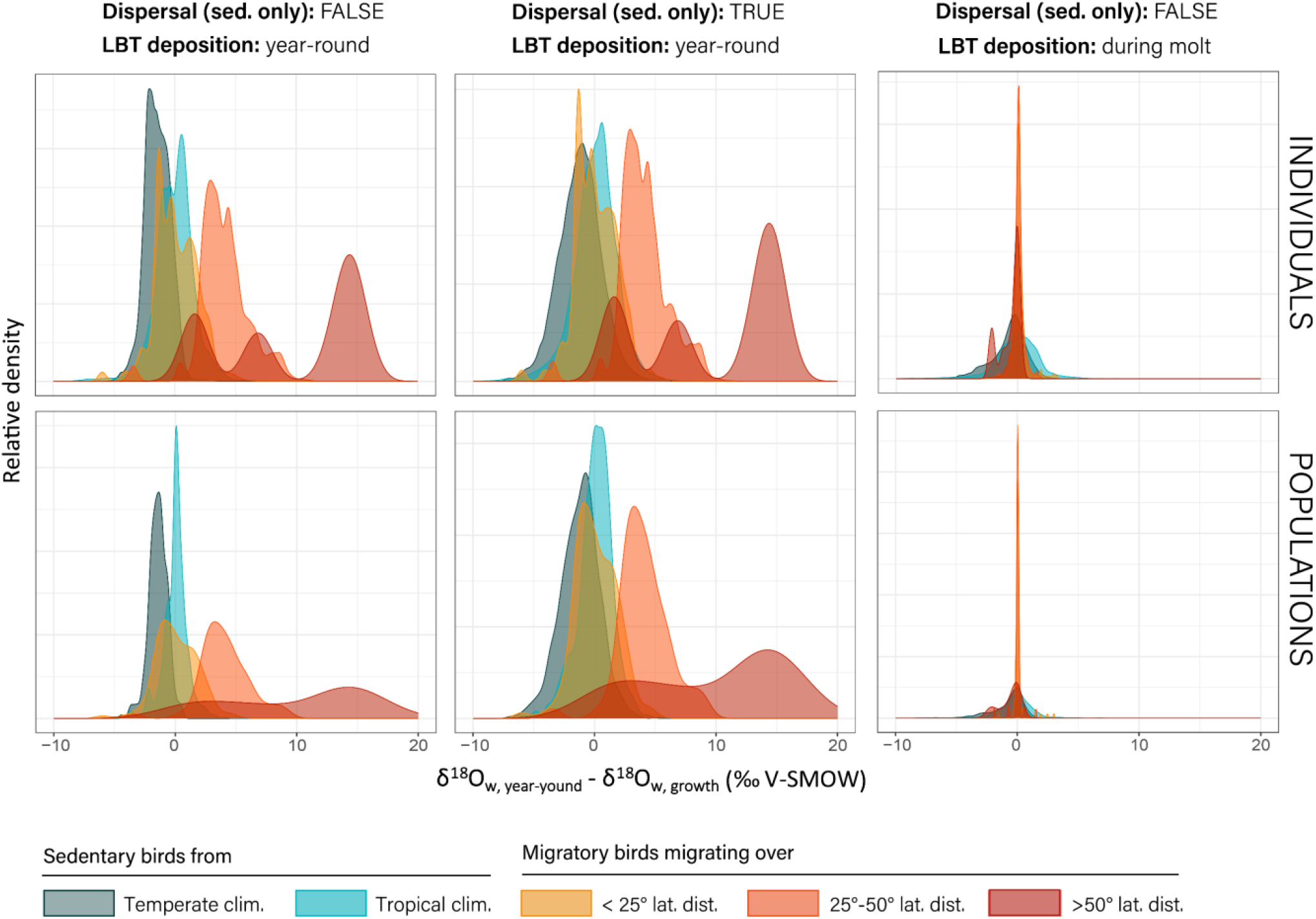
Theoretical contrasts between δ18Ow values of year-round and growth-period rainwater, under different migratory behaviour, dispersal scenarios, and LBT deposition timings (detailed in 2.2.2). Results are shown for (a) individuals and (b) populations (i.e., averaged over individuals of the same species sharing the same year-round location but differing in birth timing).

**FIGURE S9:**
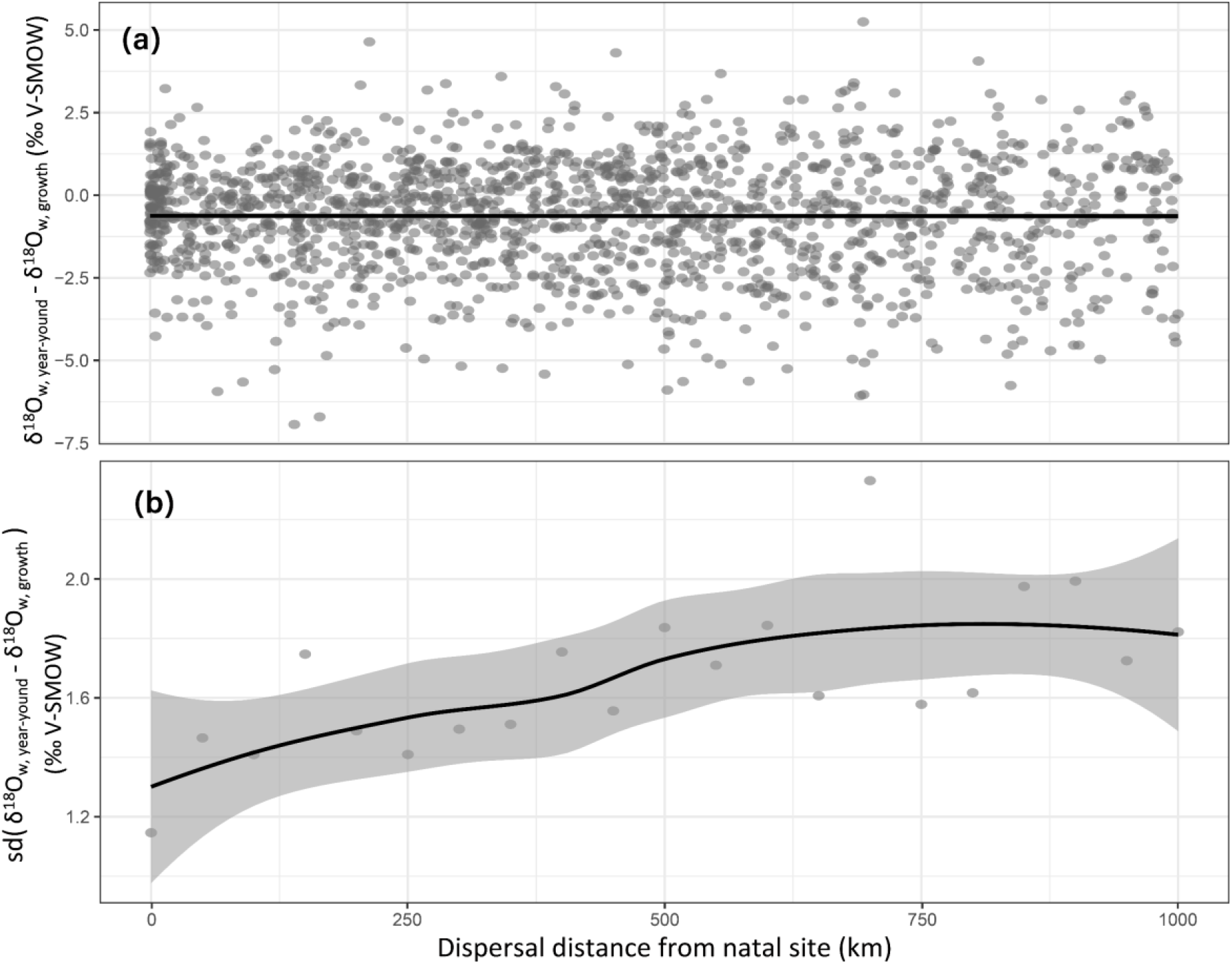
For each modelled sedentary individual (one dot = one individual), mean and standard deviation of theoretical contrasts between δ^18^O_w_ values of year-round and growth-period rainwater, depending on the dispersal distance from the natal site.

**FIGURE S10:**
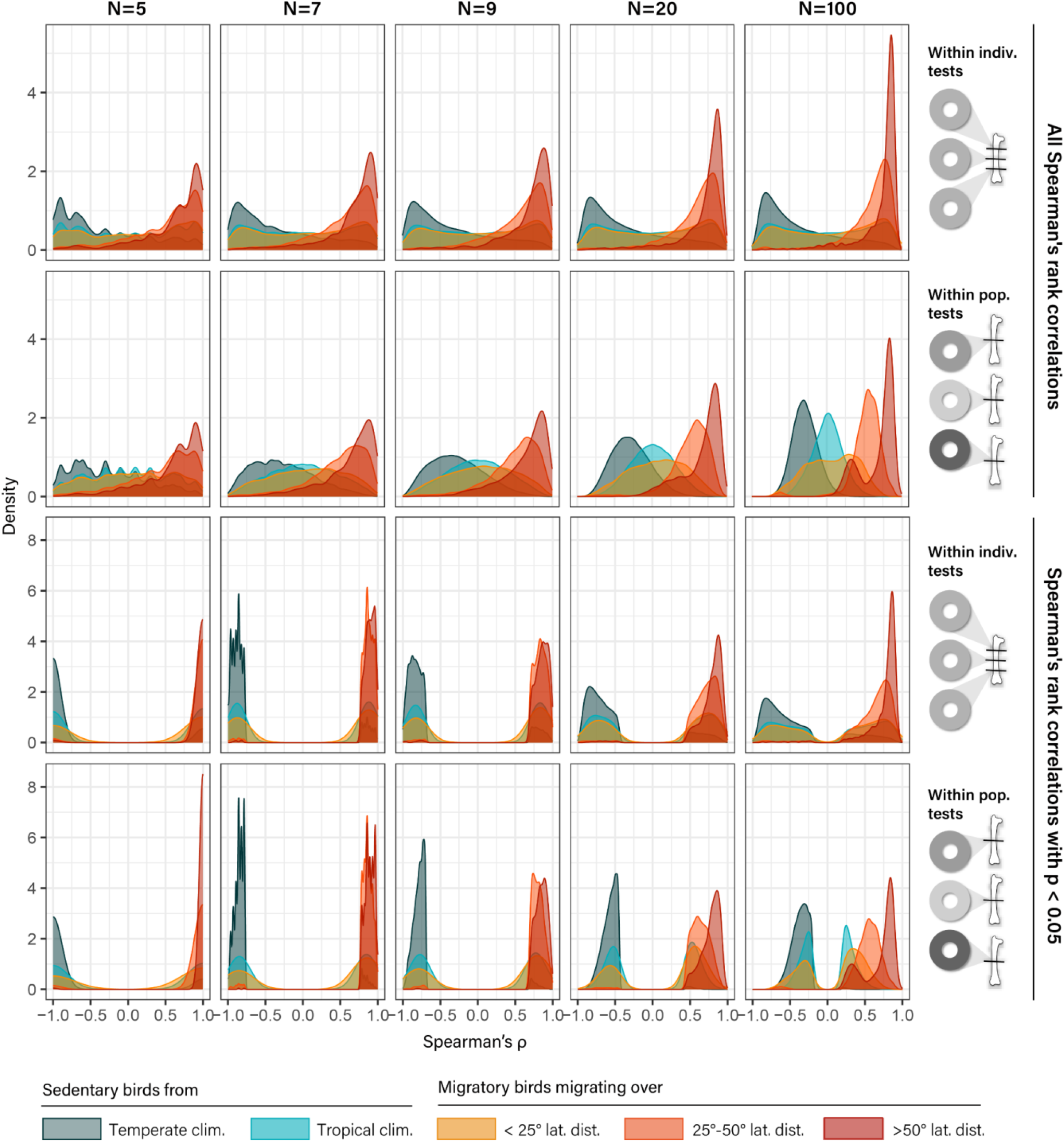
Distribution of Spearman’s *ρ* coefficients for correlations (p-value<0.05) between *f*_*LBT*_ and δ^18^O_p, bulk_ values in migratory, temperate and tropical sedentary birds. Correlations were tested on samples of varying sizes (*N*=5, 7, 9, 20 or 100), drawn either within a single individual (fixed δ^18^O_p, EBT_ values) or from different individuals within the same populations (varying δ^18^O_p, EBT_ values). All other model parameters were fixed at their default values (see Table 1 or Fig. S13). Each *N*-sized sampling was repeated 100 times per population.

**FIGURE S11:**
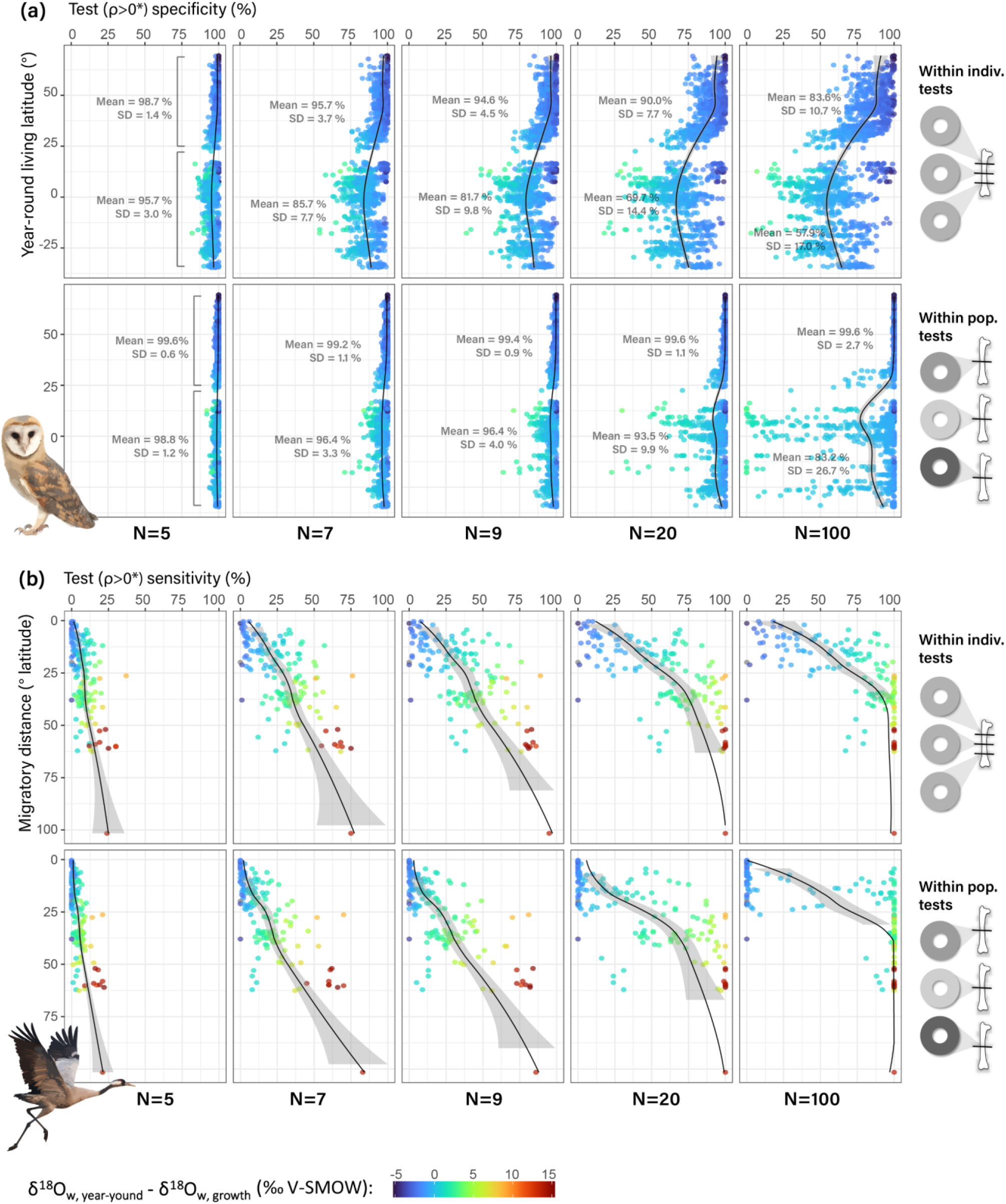
Theoretical specificity (a) and sensitivity (b) of the “Climatic Niche Contrast Test”, using the “*ρ*>0 *” decision rule: i.e., individuals or populations are classified as <u>migratory</u> (“positive” outcome) when they exhibit significantly positive correlations between *f*_*LBT*_ and δ^18^O_p, bulk_ (Spearman’s *ρ*>0 and p-value<0.05). This decision rule was applied to samples of varying sizes (*N*=5, 7, 9, 20 or 100), drawn either from repeated measurements within a single individual (fixed δ^18^O_p, EBT_ values) or from individuals within the same population (varying δ^18^O_p, EBT_ values). All other model parameters were fixed at their default values (see Table 1 or Fig. S13). Sensitivity (proportion of true positives) and specificity (1 – proportion of false positives) were computed from 100 *N*-sized samples for each individual / population. Specificity was plotted for sedentary individuals or populations against their living latitude, while sensitivity was plotted for migratory individuals / populations against their migratory distance. Observations are coloured according to the individual’s or population’s mean difference between δ^18^O values of year-round and growth-period rainwater.

**FIGURE S12:**
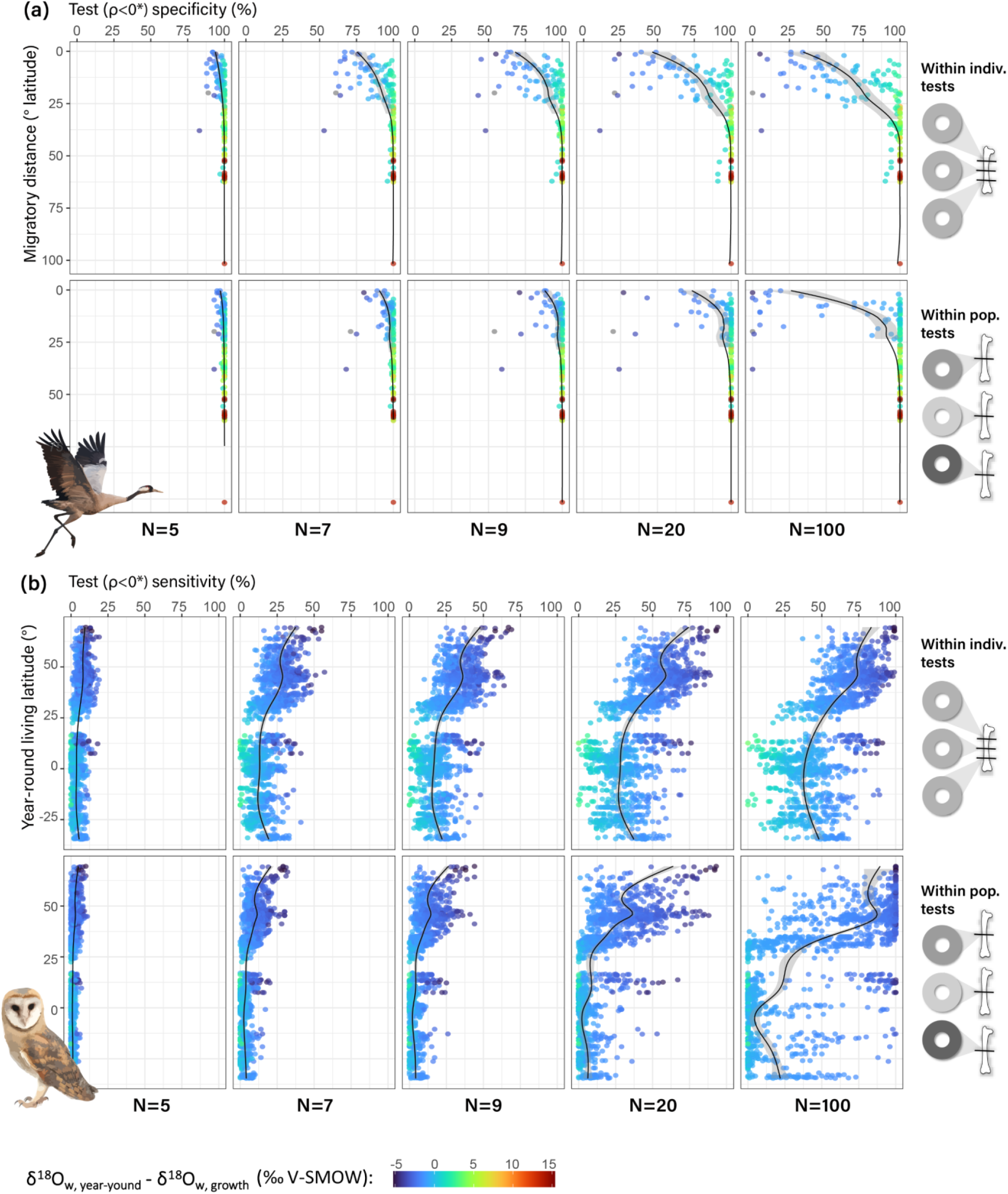
Theoretical specificity (a) and sensitivity (b) of the “Climatic Niche Contrast Test”, using the “*ρ*<0 *” decision rule: i.e., individuals or populations are classified as <u>sedentary</u> (“positive” outcome) when they exhibit significantly negative correlations between *f*_*LBT*_ and δ^18^O_p, bulk_ (Spearman’s *ρ*<0 and p-value<0.05). This decision rule was applied to samples of varying sizes (*N*=5, 7, 9, 20 or 100), drawn either from repeated measurements within a single individual (fixed δ^18^O_p, EBT_ values) or from individuals within the same population (varying δ^18^O_p, EBT_ values). All other model parameters were fixed at their default values (see Table 1 or Fig. S13). Sensitivity (proportion of true positives) and specificity (1 – proportion of false positives) were computed from 100 *N*-sized samples for each individual or population. Specificity was plotted for migratory individuals / populations against their migratory distance, while sensitivity was plotted for sedentary individuals or populations against their living latitude. Observations are coloured according to the individual’s or population’s mean difference between δ^18^O values of year-round and growth-period rainwater.

## Notes

https://doi.org/10.5281/zenodo.14825377

### Competing Interest Statement

The authors have declared no competing interest.

### Summary of Updates

This revised manuscript includes (i) an explicit mathematical formulation of the model; (ii) an additional model parameter (dispersal); (iii) a more detailed description of the model parameters and of the two statistical tests developed to infer migratory behaviour; and (iv) a sensitivity analysis evaluating how variations in key parameters influence the performance of these tests.

